# Atomistic Prediction of Structures, Conformational Ensembles and Binding Energetics for the SARS-CoV-2 Spike JN.1, KP.2 and KP.3 Variants Using AlphaFold2 and Molecular Dynamics Simulations: Mutational Profiling and Binding Free Energy Analysis Reveal Epistatic Hotspots of the ACE2 Affinity and Immune Escape

**DOI:** 10.1101/2024.07.09.602810

**Authors:** Nishank Raisinghani, Mohammed Alshahrani, Grace Gupta, Gennady Verkhivker

## Abstract

The most recent wave of SARS-CoV-2 Omicron variants descending from BA.2 and BA.2.86 exhibited improved viral growth and fitness due to convergent evolution of functional hotspots. These hotspots operate in tandem to optimize both receptor binding for effective infection and immune evasion efficiency, thereby maintaining overall viral fitness. The lack of molecular details on structure, dynamics and binding energetics of the latest FLiRT and FLuQE variants with the ACE2 receptor and antibodies provides a considerable challenge that is explored in this study. We combined AlphaFold2-based atomistic predictions of structures and conformational ensembles of the SARS-CoV-2 Spike complexes with the host receptor ACE2 for the most dominant Omicron variants JN.1, KP.1, KP.2 and KP.3 to examine the mechanisms underlying the role of convergent evolution hotspots in balancing ACE2 binding and antibody evasion. Using the ensemble-based mutational scanning of the spike protein residues and computations of binding affinities, we identified binding energy hotspots and characterized molecular basis underlying epistatic couplings between convergent mutational hotspots. The results suggested that the existence of epistatic interactions between convergent mutational sites at L455, F456, Q493 positions that enable to protect and restore ACE2 binding affinity while conferring beneficial immune escape. To examine immune escape mechanisms, we performed structure-based mutational profiling of the spike protein binding with several classes of antibodies that displayed impaired neutralization against BA.2.86, JN.1, KP.2 and KP.3. The results confirmed the experimental data that JN.1, KP.2 and KP.3 harboring the L455S and F456L mutations can significantly impair the neutralizing activity of class-1 monoclonal antibodies, while the epistatic effects mediated by F456L can facilitate the subsequent convergence of Q493E changes to rescue ACE2 binding. Structural and energetic analysis provided a rationale to the experimental results showing that BD55-5840 and BD55-5514 antibodies that bind to different binding epitopes can retain neutralizing efficacy against all examined variants BA.2.86, JN.1, KP.2 and KP.3. The results support the notion that evolution of Omicron variants may favor emergence of lineages with beneficial combinations of mutations involving mediators of epistatic couplings that control balance of high ACE2 affinity and immune evasion.

## Introduction

The Spike (S) glycoprotein of SARS-CoV-2 plays a pivotal role in the virus’s ability to enter host cells. The wealth of structural and biochemical investigations conducted on the S glycoprotein has provided crucial insights into the mechanisms that regulate virus transmission and immune evasion. The conformational flexibility of the S glycoprotein, particularly within the S1 subunit and its various domains, which includes the N-terminal domain (NTD), the receptor-binding domain (RBD), and two structurally conserved subdomains—SD1 and SD2, allows it to adapt to different stages of the viral entry process [1–9]. The transitions between closed and open states, driven by conformational changes in the NTD and RBD, enable the virus to effectively engage with host cell receptors while evading immune surveillance through structural variability [10–15]. Understanding these structural dynamics is crucial for designing therapeutics and vaccines that target the S protein, aiming to disrupt viral entry and prevent infection by SARS-CoV-2.

Biophysical investigations have delineated how thermodynamic principles and kinetic factors govern the mechanisms of the S protein [16–18]. These studies have revealed that mutations within the S protein, particularly in the S1 subunit, can induce structural alterations that affect its stability and conformational dynamics, particularly the protein’s ability to switch between the open and closed states that can influence the accessibility of the RBD critical for viral attachment to host cells. Moreover, long-range interactions between the dynamic S1 subunit (including domains like the N-terminal domain and RBD) and the more rigid S2 subunit (involved in membrane fusion) play crucial roles in determining the overall architecture and functional states of the S protein trimer [16–18].

The emergence and evolution of SARS-CoV-2 variants like BQ.1.1 and XBB.1 have raised significant interest due to their distinct characteristics, including superior growth advantages and potential immune evasion capabilities [19,20]. The XBB.1.5 subvariant has evolved through recombination events within the BA.2 lineage, incorporating genetic material from BA.2.10.1 and BA.2.75 sublineages. XBB.1.5 is equally immune evasive as XBB.1 but may have growth advantage by virtue of the higher ACE2 binding as F486P in the XBB.1.5 subvariant can restore most of the favorable hydrophobic contacts [21]. Further investigations have substantiated that the enhanced growth and increased transmissibility observed in the XBB.1.5 lineage likely stem from its preserved resistance to neutralization and improved affinity for binding to ACE2 receptors [22]. By October 2023, XBB sublineages such as XBB.1.5 and XBB.1.16, both featuring the F486P substitution, had become prevalent globally (source: https://nextstrain.org/) [23]. Compared to XBB.1.5, XBB.1.16 exhibits two substitutions: E180V in the NTD and T478R in the receptor-RBD [24]. These emerging variants demonstrated increased infectivity and transmissibility compared to earlier Omicron variants. Moreover, several residues in the RBD (R346, K356, K444, V445, G446, N450, L452, N460, F486, F490, R493, and S494) are mutated in at least five distinct new Omicron lineages. The lineage of XBB descendants, including EG.5 and EG.5.1, which carry an additional mutation F456L, has become one of the dominant lineages currently. EG.5 evolved from Omicron XBB.1.9 and bears only one additional substitution, F456L, compared to XBB.1.5. Its direct descendant, EG.5.1, features Q52H in the NTD and F456L in the RBD [25]. EG.5 and EG.5.1 were discovered to exhibit moderate resistance to antibody neutralization compared to XBB.1.5 and this resistance is particularly pronounced against class 1 monoclonal antibodies, primarily due to a single F456L mutation in the RBD [26]. Further studies specifically on the immune evasion of the EG.5.1 subvariant confirmed that the enhanced neutralization escape observed is primarily driven by the F456L mutation rather than the Q52H mutation [27]. XBB subvariants carrying both the L455F and F456L combination of flipped substitutions are often referred to as “FLip” variants and these variants were identified in over 20% of global XBB samples collected at the beginning of September 2023 [23]. The FLip variants include JG.3 (XBB.1.9.2.5.1.3.3), JF.1 (XBB.1.16.6.1), GK.3 (XBB.1.5.70.3), and JD.1.1, all of which emerged convergently. This convergence underscores that acquiring the L455F/F456L double mutation can provide a growth advantage to XBB within the human population [28].

The Omicron subvariant BA.2.86, originating from the BA.2 variant, shows substantial genetic divergence from earlier forms (Table1, Figure 1) [29–32]. Biophysical investigations have confirmed that the BA.2.86 variant can resist neutralization by monoclonal antibodies targeting epitopes in the NTD, SD1, as well as classes 1, 2, and 3 epitopes within the RBD. Notably, BA.2.86 exhibits a greater potential to evade RBD-targeted antibodies compared to the immune evasion observed in XBB.1.5 and EG.5.1 variants [29]. The immune evasion capability of the BA.2.86 subvariant was evaluated using a panel of neutralizing antibodies that are effective against XBB.1.5, revealing that BA.2.86 can evade antibodies that target XBB.1.5 [30]. Recent studies investigating the structure and binding characteristics of the BA.2.86 spike protein with ACE2 and antibodies have shown that the mutations acquired by BA.2.86 do not result in significant changes in antibody evasion compared to XBB.1.5 [32]. However, the RBD of BA.2.86 displays a 2.2-fold increase in affinity for ACE2 compared to XBB.1.5, thereby providing BA.2.86 with a transmission advantage. The latest cryo-EM structure of ACE2 complexed with BA.2.86 trimeric S protein supported the notion that the enhanced binding affinity of BA.2.86 may be driven by the electrostatic complementarity between BA.2.86 RBD and ACE2 and also potentially enhanced by the flexibility of the RBDs, allowing better exposure of the ACE2 binding within the trimer structure [32[. Another study also suggested that BA.2.86 does not have greater immune escape relative t XBB.1.5 from neutralizing immunity elicited by either Omicron XBB-family subvariant infection but improves virus fitness through greater binding affinity to ACE2 [33].

**Figure 1.**
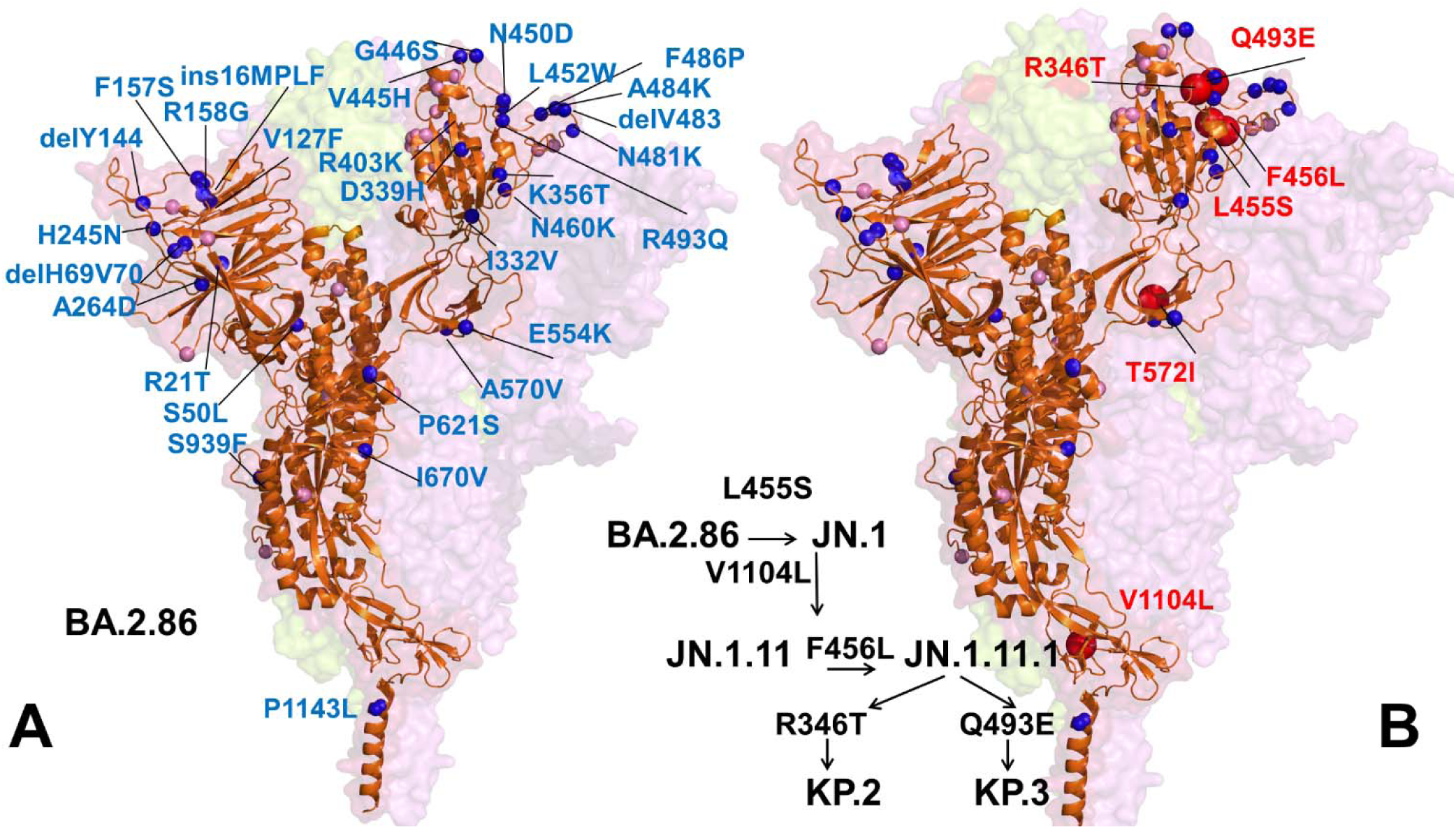
Structural overview of the SARS-CoV-2 S protein and S-RBD for Omicron BA.2.86 (A) and JN.1/KP.2/KP.3 variants (B). The S protein is shown on orange ribbons (single monomer) with the S protein trimer structure shown in surface with the reduced transparency. The BA.2.86 RBD mutations are projected onto crystallographic RBD conformation (orange ribbons) in the BA.2.86 RBD-ACE2 complex, pdb id 8QSQ. The positions of unique BA.2.86 S mutations relative to its ancestral BA.2 variant are shown in blue-colored spheres and fully annotated. BA.2 mutational positions are in pink spheres. The unique JN.1, KP.2 and KP.3 S mutations on panel (B) are shown in red spheres, and BA.2.86 mutations are in blue spheres and annotated.

JN.1 is a variant of BA.2.86 which emerged independently from Omicron BA.2 and harbors an additional L455S mutation responsible for enhanced immune escape [34]. BA.2.86/JN.1 also showed a long genetic distance from XBB.1.5 and XBB.1.16 (Table 1, Figure 1). The JN.1 variant is antigenically distinct from the XBB.1.5 variant. A comparative biochemical analysis using surface plasmon resonance (SPR) assays showed a notable reduction in ACE2 binding affinity for JN.1 indicating that its enhanced immune evasion capabilities come at the expense of reduced ACE2 binding [34]. Despite only a single additional mutation (L455S) compared to its predecessor BA.2.86, which results in increased resistance to humoral immunity, the JN.1 variant swiftly became predominant in Europe and outcompeted the previously dominant XBB lineage by early 2024 [34,35]. In vitro ACE2 binding assay showed that the dissociation constant value of the JN.1 RBD was significantly higher than that of the BA.2.86 RBD indicating that L455S mutation leads to the decreased binding affinity while it displays robust immune resistance, particularly against antibodies induced by the XBB.1.5 vaccine [35]. JN.1 demonstrated increased evasion against RBD class-1 antibodies, such as S2K146 and Omi-18, and the class-3 antibody S309 corroborating findings from other related studies [28,34].

**Table 1.**
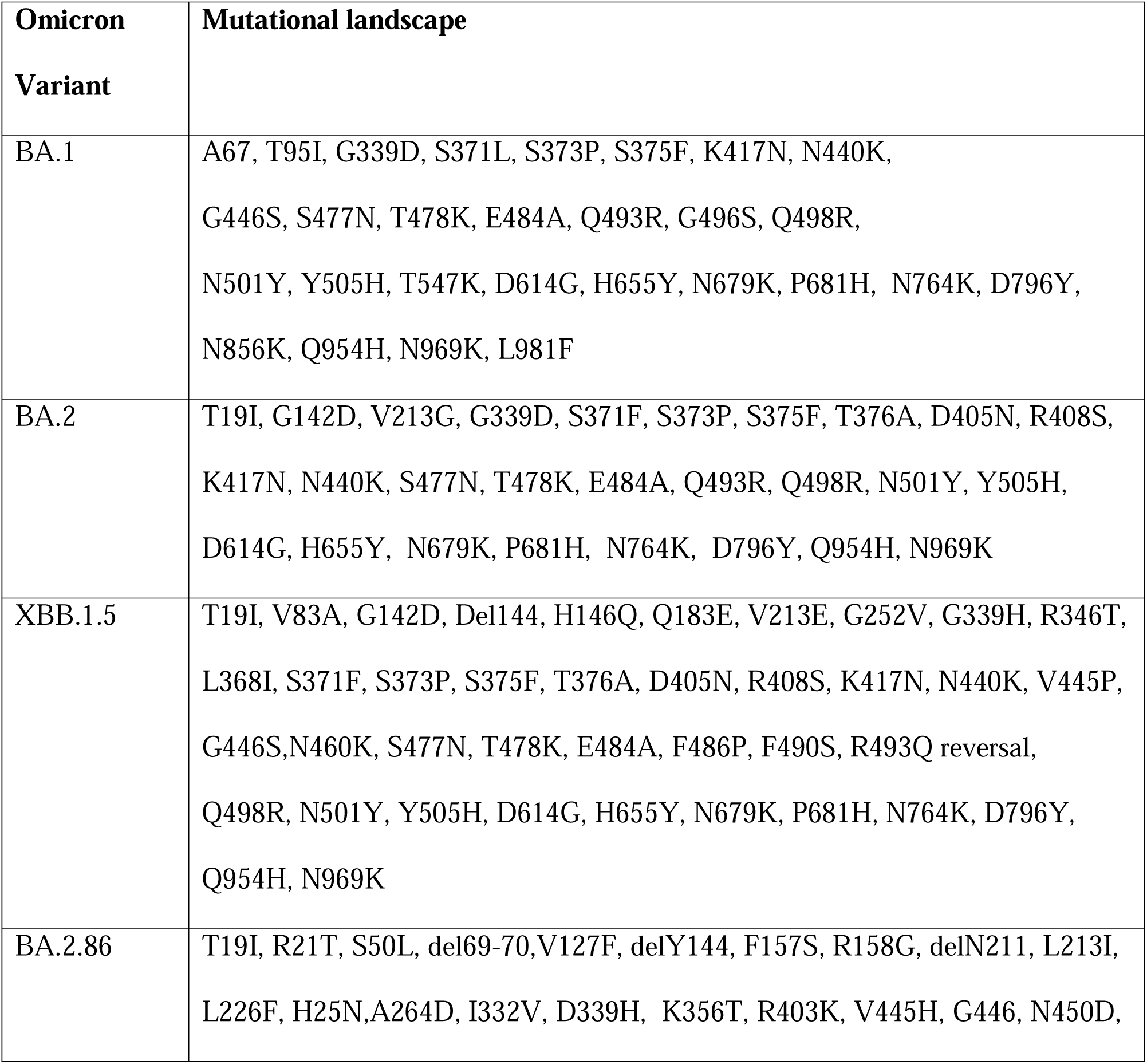
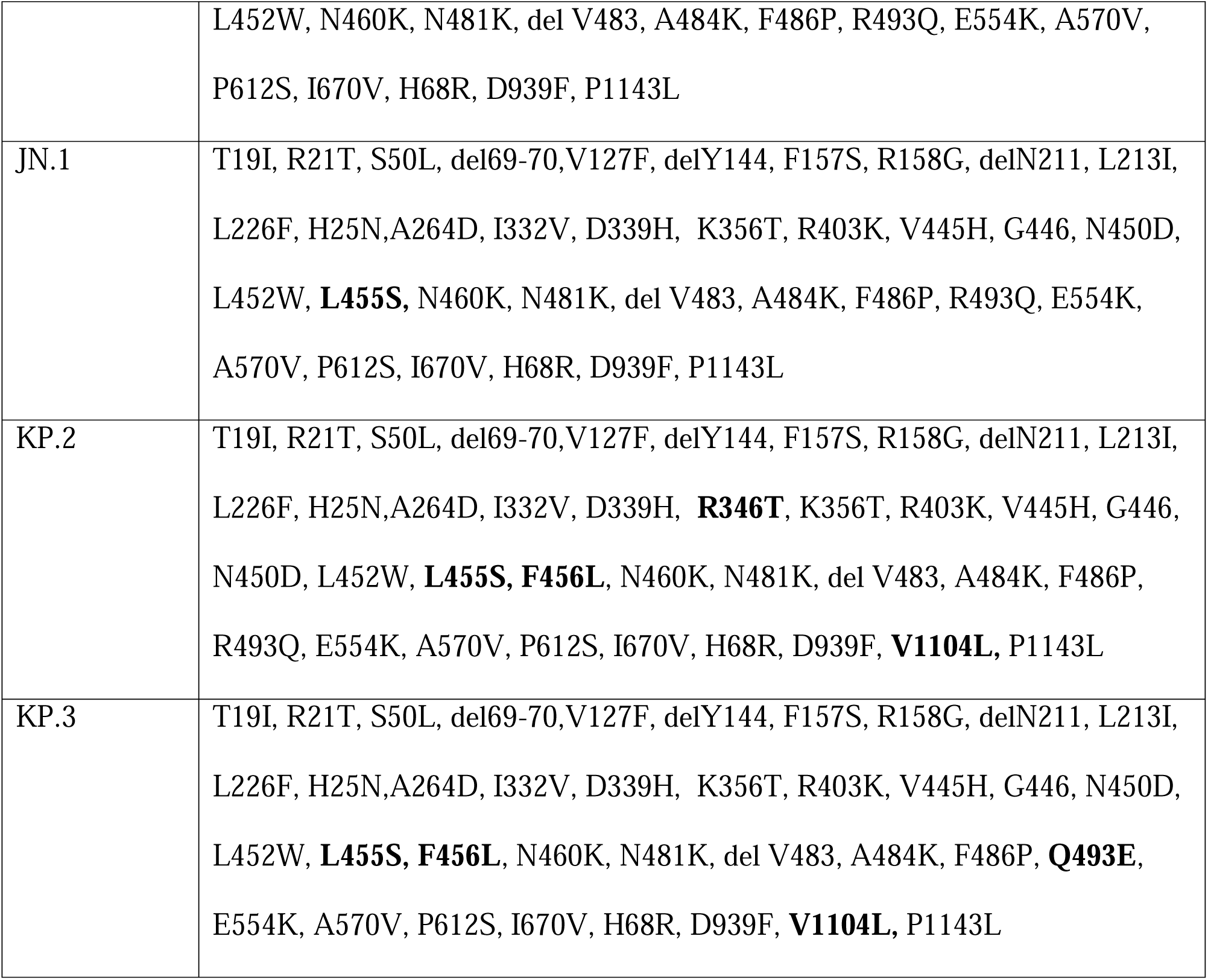
Mutational landscape of the Omicron variants.

A series of variants with mutations at L455, F456, and R346 convergent hotspot sites emerged including “SLip” variant which has the JN.1 mutations (L455S) with the additional F456L mutation [36]. More recently, we have seen the emergence of the FLiRT variant, which harbors an additional R346T mutation in the backbone of SLip. Recent studies revealed that that F456L (SLip) and R346T (FLiRT) subvariants of JN.1 contribute to further escape of JN.1-derived variants from neutralizing antibodies [36,37]. Since January 2024, JN.1 diversified into a number of sublineages, many of which share recurrent mutations R346T (JN.1.18), F456L (JN.1.16), T572I (JN.1.7), or combinations of these mutations (KP.2 variant). JN.1 subvariants including KP.2 (JN.1.11.1.2) and KP.3 (JN.1.11.1.3), which convergently acquired S protein substitutions such as S:R346T, S:F456L, and S:Q493E, and V1104L have emerged concurrently [38,39]. KP.2 and KP.3 are members of a “FLiRT” group of variants. Other FLiRT variants, including KP.1.1, have also been identified as circulating in the U.S., but have not yet become as widespread as KP.2 or KP.3 [38,39]. JN.1 subvariants with one or more of these recurrent mutations, such as KP.2 (R346T, F456L and V1104L), appear to have a growth advantage, becoming more dominant in multiple regions as of April 2024 [38].

Virological properties of KP.2 variant were examined showing that that the infectivity of KP.2 is significantly lower than that of JN.1 and that KP.2 has an increased immune resistance ability compared to JN.1 [38]. Another JN.1 descendant termed “FLuQE” variants (KP.3) continues to dominate “FLiRT” and show strong growth. KP.3 (JN.1.11.1.3), featured mutations R346T, L455S, F456L, Q493E and V1104L [39]. The pseudovirus infectivity of KP.2 and KP.3 was significantly lower than that of JN.1. Furthermore, JN.1 subvariants such as LB.1 and KP.2.3, which convergently acquired S31 deletion in addition to the above substitutions, have emerged and spread as of June 2024 and contribute to immune evasion and the increased relative effective reproduction number [40]. Importantly, LB.1 and KP.2.3 exhibited higher pseudovirus infectivity and more robust immune resistance than KP.2 showing that S31del is important for the increased infectivity and enhanced immune evasion. Both KP.2 and KP.3 variants share the F456L mutation that is critical for enhancing antibody evasion by impairing binding to class 1 RBD antibodies, confirming that antibody evasion can represent a major selective advantage for viral spread [41]. According to recent findings from Cao lab [41] and private communications of June 4 (https://twitter.com/yunlong_cao) KP.3 variant is starting to outcompete KP.2 due to Q493E mutation that enables the higher ACE2 binding affinity than KP.2. and enhanced immune evasion primarily against class I of RBD antibodies. Moreover, KP.3 (JN.1+F456L+Q493E) is the most immune evasive variant and is also the fastest-growing JN.1 sublineage. The additional F456L and Q493E mutation allows KP.3 to evade a substantial proportion of JN.1-effective antibodies, especially Class 1 antibodies.

Deep mutational scanning (DMS) experiments and functional studies suggested that evolutionary windows for the Omicron variants could be enhanced through epistatic interactions between variant mutations where the effect of one mutation can be altered depending on the presence of other mutations, resulting in non-additive impacts of mutations on specific functions [42–46]. These experiments showed evidence of compensatory epistasis in which immune escape mutations can individually reduce ACE2 binding but are compensated through epistatic couplings with affinity-enhancing Q498R and N501Y mutations [45,46]. Recent DMS experiments examined the impacts of all mutational changes and single-codon deletions in the XBB.1.5 RBDs on ACE2-binding affinity and RBD folding efficiency, revealing the expanded character of epistatic couplings between RBD residues including epistatic interactions between R493Q reversed mutations and mutations at positions Y453, L455, and F456 [47]. It was also demonstrated through mutational surveys of Omicron variants up until BA.2.86 that Q493E is typically detrimental to ACE2 binding, often incurring up to 10-fold of loss in the binding affinity [47]. In recent correspondence, Starr and colleagues reported that when Q493E mutation is combined with L455S and F456L mutations of FliRT variants, the loss of binding affinity is reversed and instead showed the increased ACE2 binding for KP.3 (https://x.com/tylernstarr/status/1800315116929560965). Moreover, Starr and colleagues reported on the results of human ACE2 binding assays on all combinations of L455S, F456L, and Q493E mutations in the BA.2.86 background, revealing the reversal of the deleterious effect of Q493E in the background of L455S+F456L The most recent reports from Cao’s lab showed that enhanced receptor ACE2 binding capabilities of the KP.3 variant may be enabled by a substantial synergistic effect of F456L and Q493E mutations. While F456L (K_D_ = 12 nM) and R346T + F456L (K_D_ = 11 nM) did not largely affect the hACE2-binding affinity of JN.1 (K_D_ = 13 nM), Q493E mutation of KP.3 substantially improved the receptor binding affinity showing K_D_ = 6.9 nM which indicates non-additive epistatic interactions between Q493E and other mutations of KP.3, particularly F456L in comparison with these pre-BA.2.86 variants. [48]. This insightful study also shows that high affinity of KP.3 due epistasis, may enable the incorporation of A475V (this mutation is convergently observed in JD.1.1, HK.3.14, JN.4, and KP.2.3.1 subvariants) for further immune evasion, given the observed only small reduction in ACE2 binding affinity for KP.3 + A475V (K_D_ is 22 nM) [48]. This study indicated that the high ACE2-binding affinity of KP.3 variant may drive the rapid transmission and prevalence of this subvariant and its descendants, enhancing its potential to acquire additional immune-evasive mutations [47]. Several other recent studies confirmed that while JN.1 displays lower affinity to ACE2 and higher immune evasion properties compared to BA.2.86, indicating that new emerging subvariants bearing Q493E mutation in the background of FLiRT variants could recover ACE2 binding affinity and continue to display the enhanced antibody evasion profile [49].

Computer simulations provided important atomistic and mechanistic advances into understanding the dynamics and function of the SARS-CoV-2 S proteins [50–55]. A series of experimental and computational studies revealed that the SARS-CoV-2 S protein can function as an allosteric regulatory machinery that is controlled by stable allosteric hotspots to modulate specific regulatory and binding functions [56–62]. Our recent studies demonstrated that convergent Omicron mutations such as G446S, F486V, F486P, F486S, and F490S can display epistatic couplings with the major stability and binding affinity hotspots which may allow for the observed broad antibody resistance [60] Analysis of conformational dynamics, binding and allosteric communications in the Omicron S protein complexes with the ACE2 host receptor characterized regions of epistatic couplings that are centered at the binding affinity hotspots N501Y and Q498R [61]. MD simulations and Markov state models systematically characterized conformational landscapes of XBB.1, XBB.1.5 Omicron variants and their complexes showing that convergent mutation sites could control evolution allosteric pockets through modulation of conformational plasticity in the flexible adaptable regions [62]. AlphaFold2-based structural modeling approaches were combined with all-atom MD simulations and mutational profiling of binding energetics and stability for prediction of dynamics, and binding of the SARS-CoV-2 Omicron BA.2.86 spike variant with ACE2 host receptor [63]. This study quantified the role of the BA.2 and BA.2.86 backgrounds in modulating binding free energy changes revealing critical variant-specific contributions of the BA.2.86 mutational sites R403K, F486P and R493Q. AlphaFold predictions of multiple conformations with MD simulations identified important differences in the conformational landscapes and binding energetics of the XBB variants and revealing the mediating role of Q493 hotspot in epistatic couplings between L455F and F456L convergent mutations [64]. The results of mutational scanning and binding analysis of the Omicron XBB spike variants with the ACE2 and a panel of class 1 antibodies provided a quantitative rationale to the experimental evidence that epistatic interactions of physically proximal binding hotspots Y501, R498, Q493, L455F and F456L residues can determine strong ACE2 binding, while convergent mutational sites F456L and F486P are instrumental in mediating broad antibody resistance [65].

We employed an integrative computational approach in which structure and conformational ensembles of the of the JN.1, KP.2 and KP.3 RBD-ACE2 complexes were first predicted using AlphaFold2 (AF2) methods [66,67] using shallow multiple sequence alignment (MSA) approach [68–70]. The Molecular Mechanics/Generalized Born Surface Area (MM-GBSA) approach is then employed for binding affinity computations of the Omicron RBD-ACE2 complexes. We combined mutational profiling of the RBD residues with MM-GBSA approach for binding affinity computations of the RBD-ACE2 complexes across subvariants BA.2.86, JN.1, BA.2.86+F456L, BA.2.86+Q493E, BA.2.86+F456L/Q493E, BA.2.86+L455S/F456L (KP.1), BA.2.86+L455S/Q493E, KP.2 and BA.2.86+L455S/F456L/Q493E (KP.3). Using mutational profiling of the S residues we identify binding energy hotspots and quantify epistatic couplings between convergent mutational hotspots. We examine a hypothesis that the emerging new variants may induce epistasis patterns where structural stability of the RBD can promote evolvability in vulnerable regions by tolerating combinations of convergent mutations at L455, F456, Q493 positions that confer beneficial phenotypes. Consistent with the latest functional studies of FLiRT and FLuQE variants, we found that that the Q493E mutation may be epistatically coupled with the F456L mutation, resulting in the reversal of the detrimental effect of Q493E seen in other backgrounds. The results showed that combining the AF2 predictions conformational ensembles of RBD-ACE2 complexes with MM-GBSA computations of the RBD-ACE2 binding can produce robust quantitative analysis of binding mechanisms. In addition, we also conducted mutational scanning of the S complexes with several important class I monoclonal antibodies to quantify the effect of convergent Omicron mutations on immune escape and the mechanism underlying balance between ACE2 binding and immune evasion of BA.2.86, JN.1, KP.2 and KP.3 variants. The panel of studied antibodies included RBD-targeting antibodies such as S2K146 [71], Omi-3[72] Omi-18[72], Omi-42 [72] as well as antibodies BD55-5514 (SA55) and BD55-5840 (SA58) [73,74] that bind to a different RBD epitope and are experimentally known to tolerate escape mutations in BA.2.8, JN.1, KP.2 and KP.3 variants. Structural analyses of BD-508, BD-236, BD-629, BD-604, and BD-515 that are class 1 antibodies and displayed a similar RBD-binding pose showed diverse interactions between antibodies and RBD and pointed to high diversity that RBD antibodies may exhibit [75]. We show that a group of convergent mutational sites Y453, L455 and F456 represent prevalent escape hotspots against class I antibodies as all modifications in these positions, particularly L455S and F456L, lead to a dramatic loss of binding interactions with the antibodies. A particular class of antibodies, F2 and F3 antibodies compete with ACE2, and their binding is affected by T376, K378, D405, R408 and G504, corresponding to class 1/4 which includes BD55-5514 (also known as SA55) [76]. Here, we demonstrate that SA55 and SA58 antibodies avoid targeting convergent mutational sites and therefore can overcome immune evasion. The results suggest a mechanism in which convergent Omicron mutations can promote high transmissibility and antigenicity of the virus by controlling the interplay between the RBD stability and conformational adaptability, allowing for optimal fitness tradeoffs between binding to the host receptor and robust immune evasion profile.

## 2. Materials and Methods

### 2.1 AI-based structural modeling and statistical assessment of AF2 models

Structural prediction of the RBD-ACE2 complexes for JN.1, KP.2 and KP.3 variants were carried out using AF2 framework [66,67] within the ColabFold implementation [77] using a range of MSA depths and other parameters. The default MSAs are subsampled randomly to obtain shallow MSAs containing as few as five sequences. We used *max_msa* field to set two AF2 parameters in the following format: *max_seqs:extra_seqs*. Both of these parameters determine the number of sequences subsampled from the MSA (*max_seqs* sets the number of sequences passed to the row/column attention track and *extra_seqs* the number of sequences additionally processed by the main evoformer stack). The lower values encourage more diverse predictions but increase the number of misfolded models. Similar to previous studies showing that MSA depth adaptations may facilitate conformational sampling [68–70]. MSA depth was modified by setting the AF2 config.py parameters *max_extra_msa* and *max_msa_clusters* to 32 and 16, respectively. We additionally manipulated the *num_seeds* and the *num_recycles* parameters to produce more diverse outputs. We use *max_msa*: 16:32, *num_seeds*: 4, and *num_recycles*: 12. AF2 makes predictions using 5 models pretrained with different parameters, and consequently with different weights. To generate more data, we set the number of recycles to 12, which produces 14 structures for each model starting from recycle 0 to recycle 12 and generating a final refined structure. Recycling is an iterative refinement process, with each recycled structure getting more precise. Each of the AF2 models generates 14 structures, amounting to 70 structures in total.

AF2 models were ranked by predicted Local Distance Difference Test (pLDDT) scores (a per-residue estimate of the prediction confidence on a scale from 0 to 100), quantified by the fraction of predicted Cα distances that lie within their expected intervals. The values correspond to the model’s predicted scores based on the lDDT-Cα metric, a local superposition-free score to assess the atomic displacements of the residues in the model [66,67]. Structural models were compared to the just released structures of the RBD-ACE2 complexes for BA.2.86 variant (pdb id 8QSQ) [32] using structural alignment as implemented in TM-align [78]. An optimal superposition of the two structures is then built and TM-score is reported as the measure of overall accuracy of prediction for the models. We also used the root mean square deviation (RMSD) superposition of backbone atoms calculated using ProFit (http://www.bioinf.org.uk/software/profit/).

### 2.2. Molecular Dynamics Simulations

The crystal and cryo-EM structures of the Omicron RBD-ACE2 complexes are obtained from the Protein Data Bank [79]. For simulated structures, hydrogen atoms and missing residues were initially added and assigned according to the WHATIF program web interface [80]. The missing regions are reconstructed and optimized using template-based loop prediction approach ArchPRED [81]. The side chain rotamers were refined and optimized by SCWRL4 tool [82]. The protonation states for all the titratable residues of the ACE2 and RBD proteins were predicted at pH 7.0 using Propka 3.1 software and web server [83,84]. The protein structures were then optimized using atomic-level energy minimization with composite physics and knowledge-based force fields implemented in the 3Drefine method [85,86]. We considered glycans that were resolved in the structures. NAMD 2.13-multicore-CUDA package [87] with CHARMM36 force field [88] was employed to perform 1µs all-atom MD simulations for the Omicron RBD-ACE2 complexes. The structures of the 2 S-RBD complexes were prepared in Visual Molecular Dynamics (VMD 1.9.3) [89] and with the CHARMM-GUI web server [90,91] using the Solutions Builder tool. Hydrogen atoms were modeled onto the structures prior to solvation with TIP3P water molecules [92] in a periodic box that extended 10 Å beyond any protein atom in the system. To neutralize the biological system before the simulation, Na^+^ and Cl^−^ ions were added in physiological concentrations to achieve charge neutrality, and a salt concentration of 150 mM of NaCl was used to mimic a physiological concentration. All Na^+^ and Cl^−^ ions were placed at least 8 Å away from any protein atoms and from each other. MD simulations are typically performed in an aqueous environment in which the number of ions remains fixed for the duration of the simulation, with a minimally neutralizing ion environment or salt pairs to match the macroscopic salt concentration [93]. All protein systems were subjected to a minimization protocol consisting of two stages. First, minimization was performed for 100,000 steps with all the hydrogen-containing bonds constrained and the protein atoms fixed. In the second stage, minimization was performed for 50,000 steps with all the protein backbone atoms fixed and for an additional 10,000 steps with no fixed atoms. After minimization, the protein systems were equilibrated in steps by gradually increasing the system temperature in steps of 20 K, increasing from 10 K to 310 K, and at each step, a 1ns equilibration was performed, maintaining a restraint of 10 kcal mol^−1^ Å^−2^ on the protein C atoms. After the restraints on the protein atoms were removed, the system was equilibrated for an additional 10 ns. Long-range, non-bonded van der Waals interactions were computed using an atom-based cutoff of 12 Å, with the switching function beginning at 10 Å and reaching zero at 14 Å. The SHAKE method was used to constrain all the bonds associated with hydrogen atoms. The simulations were run using a leap-frog integrator with a 2 fs integration time step. The ShakeH algorithm in NAMD was applied for the water molecule constraints. The long-range electrostatic interactions were calculated using the particle mesh Ewald method [94] with a cut-off of 1.0 nm and a fourth-order (cubic) interpolation. The simulations were performed under an NPT ensemble with a Langevin thermostat and a Nosé–Hoover Langevin piston at 310 K and 1 atm. The damping coefficient (gamma) of the Langevin thermostat was 1/ps. In NAMD, the Nosé– Hoover Langevin piston method is a combination of the Nosé–Hoover constant pressure method [95] and piston fluctuation control implemented using Langevin dynamics [96,97]. An NPT production simulation was run on equilibrated structures for 1µs keeping the temperature at 310 K and a constant pressure (1 atm).

### 2.3. Distance Fluctuations Stability and Communication Analysis

We employed distance fluctuation analysis of the simulation trajectories to compute residue-based rigidity/flexibility profiles. The fluctuations of the mean distance between each pseudo-atom belonging to a given amino acid and the pseudo-atoms belonging to the remaining protein residues were computed. The fluctuations of the mean distance between a given residue and all other residues in the ensemble were converted into distance fluctuation stability indexes that measure the energy cost of the residue deformation during simulations [98–100]. The distance fluctuation stability index for each residue is calculated by averaging the distances between the residues over the simulation trajectory using the following expression:

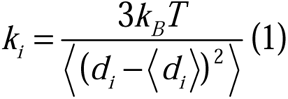

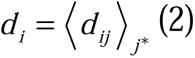

is the instantaneous distance between residue *i* and residue *j*, *k_B_* is the Boltzmann constant, *T* =300K. ⟨ ⟩ denotes an average taken over the MD simulation trajectory and *d_i_* = ⟨*d_ij_*⟩*_j*_* is the average distance from residue *i* to all other atoms *j* in the protein (the sum over *j*_*_ implies the exclusion of the atoms that belong to the residue *i*). The distances between residue *i* and residue *j* are calculated for each conformation along MD trajectories and the mean values of the inter-residue distances are obtained from averaging over the complete ensemble derived from MD simulations. The interactions between the *C_α_* atom of residue *i* and the *C_α_* atom of the neighboring residues *i*-1 and *i* +1 are excluded in the calculation since the corresponding distances are constant. The inverse of these fluctuations yields an effective force constant *k_i_* that describes the ease of moving an atom with respect to the protein structure.

### 2.4. Binding Free Energy Computations: Mutational Scanning and Sensitivity Analysis

We conducted mutational scanning analysis of the binding epitope residues for the SARS-CoV-2 S RBD-ACE2 complexes. Each binding epitope residue was systematically mutated using all substitutions and corresponding protein stability and binding free energy changes were computed. BeAtMuSiC approach [101–103] was employed that is based on statistical potentials describing the pairwise inter-residue distances, backbone torsion angles and solvent accessibilities, and considers the effect of the mutation on the strength of the interactions at the interface and on the overall stability of the complex. The binding free energy of protein-protein complex can be expressed as the difference in the folding free energy of the complex and folding free energies of the two protein binding partners:

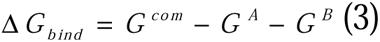

The change of the binding energy due to a mutation was calculated then as the following:

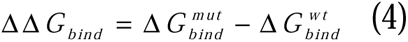

We leveraged rapid calculations based on statistical potentials to compute the ensemble-averaged binding free energy changes using equilibrium samples from simulation trajectories. The binding free energy changes were obtained by averaging the results over 1,000 and 10, 000 equilibrium samples for each of the studied systems.

### 2.5 Binding Free Energy Computations

We compute the ensemble-averaged binding free energy changes using equilibrium samples from simulation trajectories. The binding free energy changes were computed by averaging the results over 1,000 equilibrium samples for each of the studied systems. The binding free energies were initially computed for the Omicron RBD-ACE2 complexes using the MM-GBSA approach [104,105]. We also evaluated the decomposition energy to assess the energy contribution of each amino acid during the binding of RBD to ACE2 [106,107]. The binding free energy for the each RBD–ACE2 complex was obtained using:

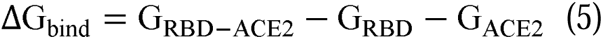

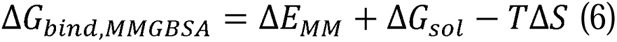

where Δ*E_MM_* is total gas phase energy (sum of Δ*E_internal_*, Δ*E_electrostatic_*, and Δ*Evdw*); Δ*Gsol* is sum of polar (Δ*G_GB_*) and non-polar (Δ*G_SA_*) contributions to solvation. Here, G_RBD–ACE2_ represent the average over the snapshots of a single trajectory of the MD RBD–ACE2complex, G_RBD_ and G_ACE2_ corresponds to the free energy of RBD and ACE2 protein, respectively. MM-GBSA is employed to predict the binding free energy and decompose the free energy contributions to the binding free energy of a protein–protein complex on per-residue basis [107]. The binding free energy with MM-GBSA was computed by averaging the results of computations over 1,000 samples from the equilibrium ensembles. The entropy contribution was not included in the calculations because the entropic differences between for estimates of binding affinities are exceedingly small owing to small mutational changes and preservation of the conformational dynamics [108,109].

## 3. Results

### 3.1. Evolutionary and Phylogenetic Analysis of Differences between XBB, BA.2.86 and FLiRT Lineages

The evolutionary differences and divergence of XBB and BA.2.86 linages among the Omicron variants are illustrated by the phylogenetic analysis using their corresponding clades nomenclature from Nextstrain an open-source project for real time tracking of evolving pathogen populations (https://nextstrain.org/). [23]. Nextstrain provides dynamic and interactive visualizations of the phylogenetic tree of SARS-CoV-2, allowing users to explore the evolutionary relationships between different lineages and variants. This approach assigns SARS-CoV-2 variant as clade when it reaches a frequency of 20% globally at any time point. A new clade should be at least 2 mutations away from its parent major clade. According to the Nextstrain evolutionary analysis (Figures 2,3) XBB.1.5 (23A clade) is a recombinant variant as it descends from XBB (22F clade). XBB.1.5 has additional S mutations S:G252V and S:S486P that are also shared in XBB.1.5+F456L (EG.5) and XBB.1.5+L455F/F456L (XBB.1.5.70). The evolutionary analysis of different Omicron lineages indicated that BA.2.86 represents the step-change evolution that has given the virus a global growth advantage beyond the XBB.1-based lineages (Figures 2,3). BA.2.86 S carries a novel and distinct constellation of mutations that produced a novel branch in the SARS-CoV-2 spike phylogenetic tree (Figures 2,3).

**Figure 2.**
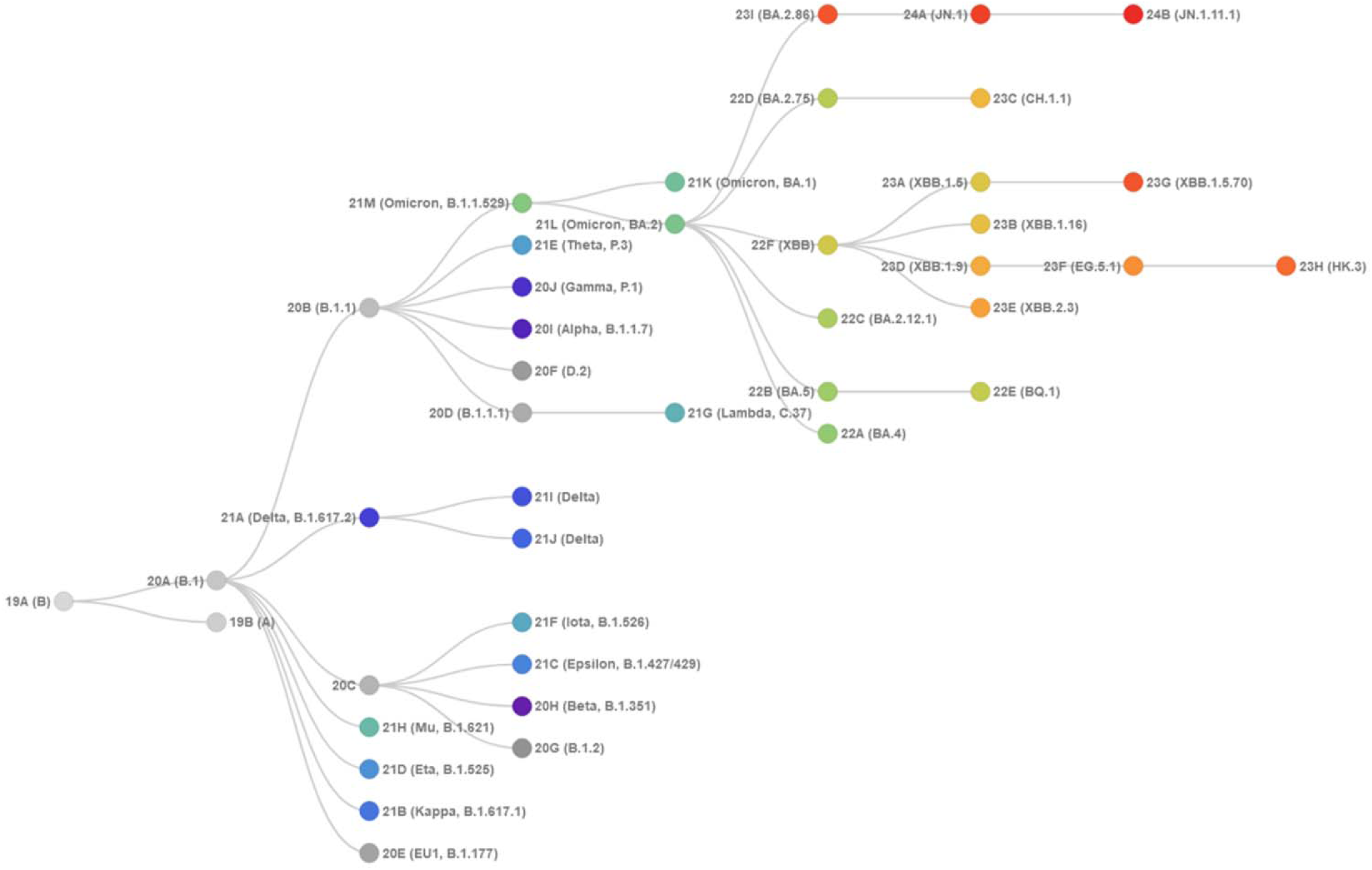
The evolutionary tree of current SARS-CoV-2 clades. XBB.1, XBB.1.5, BA.2.86, JN.1, jN.1.11.1 variants are shown on the current tree. The graph is generated using Nextstrain, an open-source project for real time tracking of evolving pathogen populations (https://nextstrain.org/) [39]. The clade 22F corresponds to XBB, 23A corresponds to XBB.1.5, 23I corresponds to BA.2.86, JN.1 corresponds to 24A and JN.1.11.1 corresponds to 24B clades. The evolutionary tree highlights divergence of BA.2.86 and JN.1 variants from XBB.1.5 lineage.

**Figure 3.**
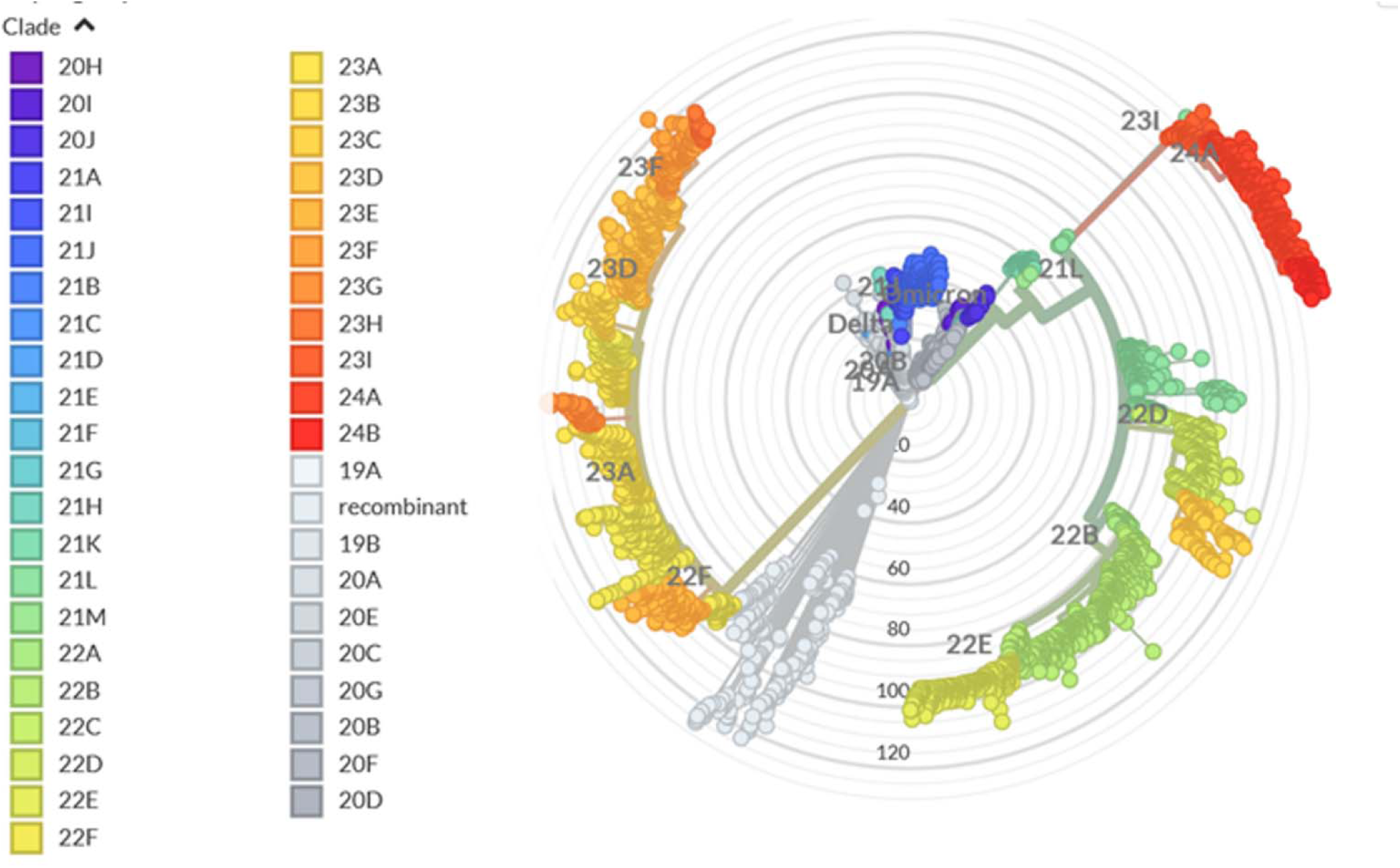
An overview of the phylogenetic analysis and divergence of Omicron variants. The graph is generated using Nextstrain, an open-source project for real time tracking of evolving pathogen populations (https://nextstrain.org/).

Compared with its BA.2 ancestor, BA.2.86 contains 34 mutations relative to BA.2 (29 substitutions, 4 deletions and 1 insertion) including RBD mutations I332V, D339H, K356T, R403K, V445H, G446S, N450D, L452W, N460K, N481K, delV483, A484K, F486P and R493Q (Figure 1). Many of these mutations such as G446S, N460K, F486P, and R493Q have been previously observed in other variants. BA.2.86 consists of eight different sublineages, namely BA.2.86, BA.2.86.1, BA.2.86.2, BA.2.86.3, JQ.1, JN.3, JN.2 and JN.1 [33,49,110].

JN.1 is a major descendent lineage of BA.2.86. JN.1 received Nextstrain clade classification 24A (Figures 2,3). In 2024 JN.1 diversified into a number of sublineages, many of which share recurrent mutations R346T (JN.1.18), F456L (JN.1.16), T572I (JN.1.7), or combinations of these mutations (KP.2 variant). These mutations can individually contribute to the enhanced antibody evasion (R346T and F456L), ACE2-binding (R346T), and S protein stability and dynamic profile (T572I) when found on the genetic background of earlier Omicron subvariants. The evolutionary diagrams show distinct evolutionary trajectories of SARS-CoV-2 Omicron XBB and BA.2.86/JN.1 lineages (Figure 3). The current evolutionary divergences between XBB and BA.2.6, JN.1 lineages illustrated by Nextstrain diagrams (Figure 3) indicate that evolutionary trajectories of Omicron lineages can proceed through diverse mechanisms including complex recombination, antigenic drift and convergent evolution that led to convergent immune-escape mutations as many lineages independently acquired mutations such as R346T, L455F/S, F456L and T572I.

### 3.2 AF2-Based Modeling and Prediction of the BA.2.86, JN.1, KP.2 and KP.3 RBD-ACE2 Complexes and Conformational Ensembles

The lack of molecular details on structure, dynamics and binding energetics of the FLiRT and FLuQE variants including primarily JN.1, KP.2 and KP.3 RBD binding with the ACE2 receptor and antibodies provides a considerable challenge that needs to be addressed to rationalize the experimental data and establish the atomistic basis for the proposed molecular mechanisms. These important objectives are addressed in the current study by using a combination of AI-based structural modeling approaches, MD simulations and the ensemble-based mutational profiling of the Omicron JN.1, KP.2 and KP.3 variants with the ACE2 receptor and a panel of monoclonal antibodies.

We fist proceeded with AF2 structural predictions of the structures and conformational ensembles of the RBD-ACE2 complexes for the BA.2.86, JN.1, KP.2 and KP.3 variants using AF2 methodology [66,67] with varied MSA depth [68–70]. By using two AF2 parameters in the following format: *max_seqs:extra_seqs* we can manipulate the number of sequences subsampled from the MSA (*max_seqs* sets the number of sequences passed to the row/column attention track and *extra_seqs* the number of sequences additionally processed by the main evoformer stack). More diverse predictions are generally encouraged with the lower values and MSA depth adaptations may facilitate conformational sampling [68–70]. The predicted structural ensembles for the BA.2.86, JN.1, KP.2 and KP.3 variants revealed a considerable heterogeneity of the RBD in all variants, showing a progressive increase in the mobility of the RBM region in KP.2 and KP.3 (Supporting Information, Figure S1). Interestingly, for BA.2.86 and JN.1 variants, structural variations in the RBD loop 444-452 are rather moderate but become more diffuse in the KP.2 and KP.3 variants (Supporting Information, Figure S1). The RBD loop regions 444-452 and 475-487 harbor BA.2.86 mutations V445H, G446S, N450D, L452W, delV483, A484K and F486P. These regions are also in the immediate structural proximity of L455, F456 and Q493 positions that undergo mutational changes in JN.1, KP.2 and KP.3 variants.

The generated RBD conformation were evaluated using the pLDDT metric. The distributions of pLDDT values displayed pronounced peaks at pLDDT ∼70-85 for all studied variants (Figure 4). The pLDDT assessments of the structural ensembles for BA.2.86 variant (Figure 4A) showed that the dominant fraction of the ensemble (pLDDT ∼80-85) conforms very closely with the experimental structure, while a minor fraction of the ensemble samples pLDDT values ∼60-80 associated with the native RBD conformations but featuring the increased mobility of the RBD flexible loops (Figure 4A). Although other pLDDT distributions were generally similar, we noticed more shallow profiles for JN.1 and KP.2 variants (Figure 4B,C). A fairly significant number of smaller peaks at various pLDDT values (was seen for the KP.3 variant with a dominant distribution peak at pLDDT ∼70-75 (Figure 4D).

**Figure 4.**
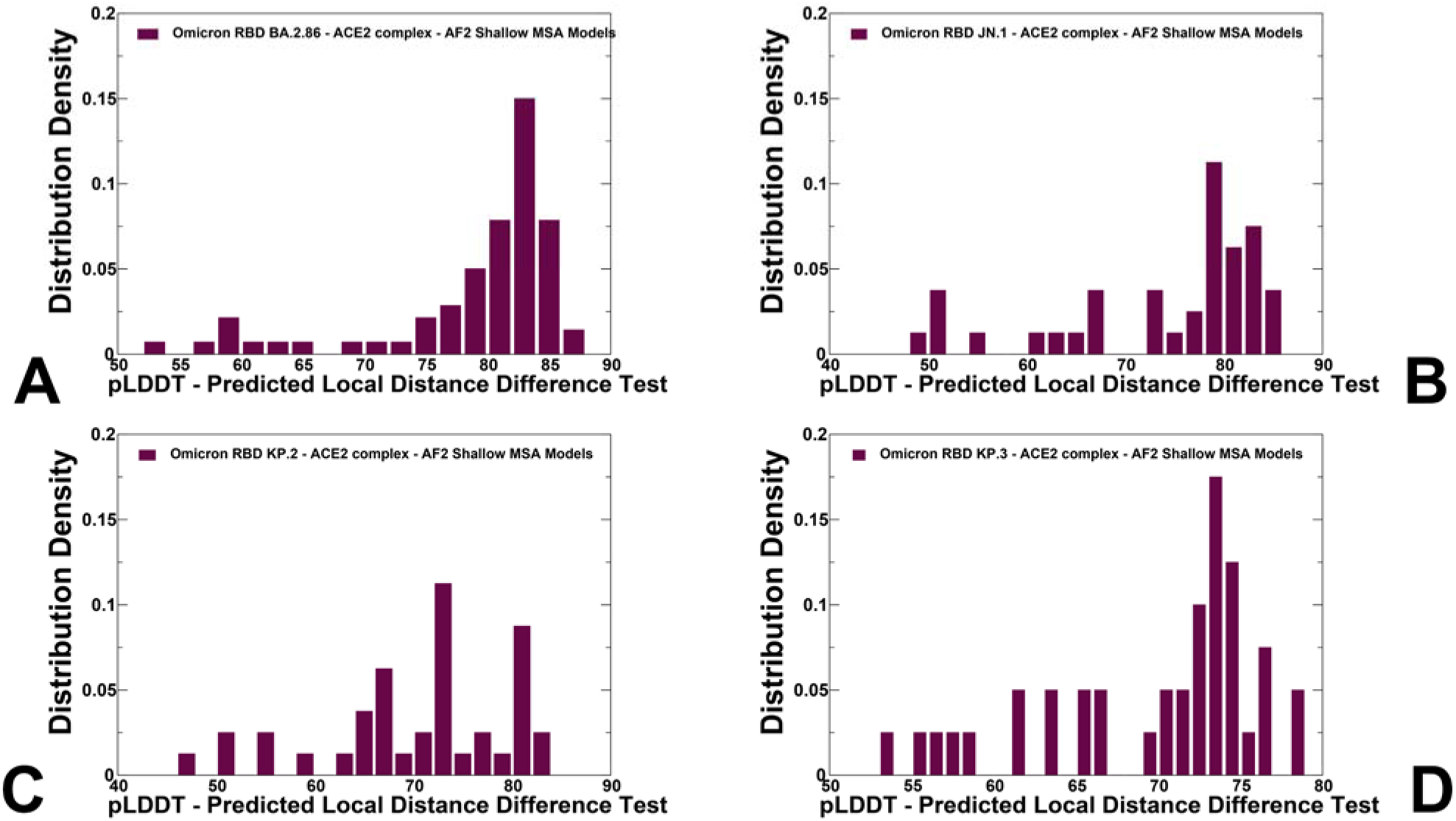
The distributions of pLDDT structural model assessment for the BA.2.86, JN.1, KP.2 and KP.3 RBD conformational ensemble obtained from AF2-MSA depth predictions. The density distribution of the pLDDT structural model estimate of the prediction confidence for BA.2.86 (A) JN.1 (B), KP.2 (C), and KP.3 (D).

AF2 predictions with pLDDT values ∼70-90 are typically associated with high confidence while the regions with pLDDT values ∼ 50-70 could indicate the reduced confidence [66,67]. Overall, the obtained conformational ensembles suggested the increased heterogeneity in the JN.1, KP.2 and especially KP.3 RBD variants which may potentially enable these variants to leverage a more mobile RBD structure to modulate and evade antibody neutralization. The distribution of structural similarity metric RMSD computed with respect to the cryo-EM structure of the BA.2.86 RBD-ACE2 complex echoed the pLDDT profile, showing strong peaks at low RMSD values ∼ 1-2.5 Å for BA.2.86 (Figure 5A) and JN.1 variants (Figure 5B). Although the profiles showed low RMSD peaks for KP.2 and KP.3 variants, these distributions also populated large RMSD values indicating the tendency of the AF2 predictions to produce overly heterogeneous conformations that could deviate significantly from the native structure of the BA.2.86 RBD (Figure 5C,D).

**Figure 5.**
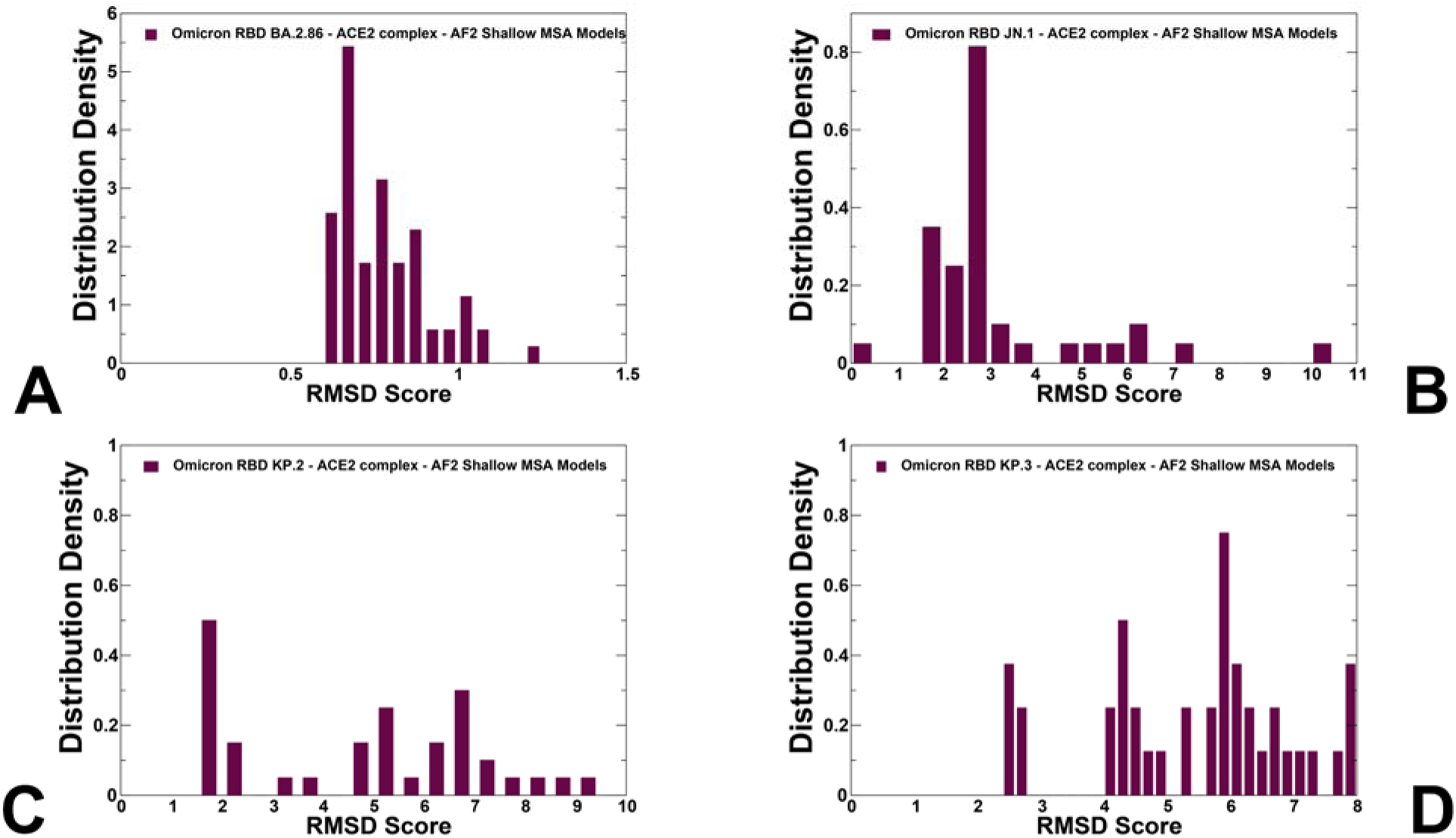
The density distribution of the RMSD score measuring structural similarity of the predicted RBD conformational ensembles with respect to the structure of the BA.2.86 RBD-ACE2 (pdb id 8QSQ) for the BA.2.86 (A) JN.1 (B), KP.2 (C), and KP.3 variants (D).

To select functionally relevant conformations from the ensemble, the predicted top models for the RBD-ACE2 complexes were selected solely based on the confidence metric pLDDT by choosing conformations with pLDDT > 70 (Figure 6). It is assumed that that conformations with high pLDDT values could be considered as functionally relevant representatives of the RBD mobility. Structural alignment of the AF2 models with the recently released cryo-EM structure of the ancestor BA.2.86 RBD-ACE2 complex (pdb id 8QSQ) yielded the RMSD < 0.8 - 1.0 Å mostly displaying moderate deviations in the RBD loops which are especially apparent in the RBM region bearing mutations F486P and N481K (Figure 6).

**Figure 6.**
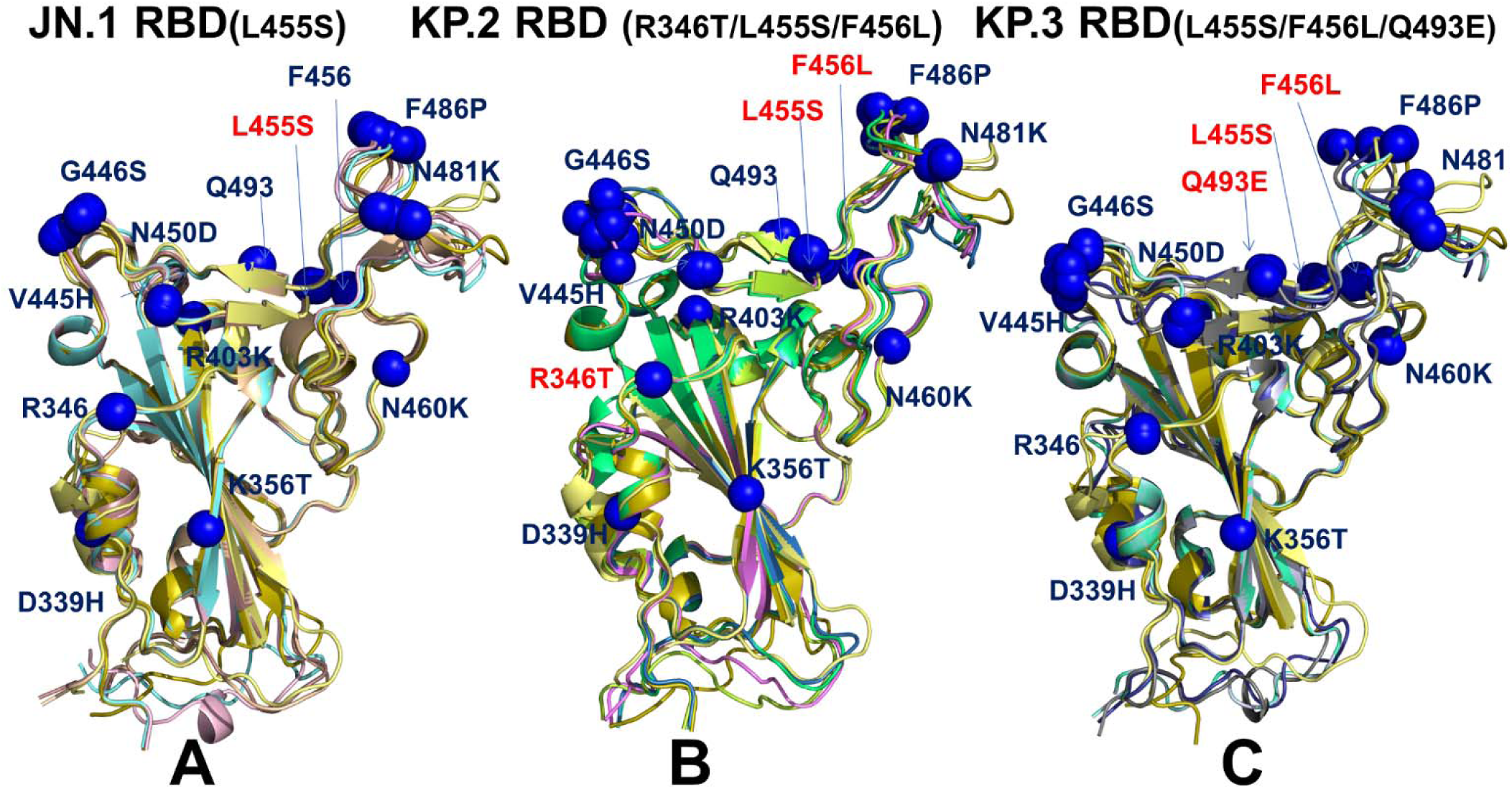
Structural alignment of the AF2-predicted RBD conformational ensembles in the complexes with ACE2 for JN.1 (A), KP.2 (B) and KP.3 variants (C). Structural alignment of the AF2-predicted conformations with high pLDDT values > 80.0 and the cryo-EM structure of the BA.2.86 RBD-ACE2 complex (pdb id 8QSQ). The RBD conformations are shown in ribbons. The sites of unique BA.2.86 mutations D339H, K356T, R403K, V445H, G446S, N450D, L452W, N460K, N481K, A484K, F486P, R493Q are shown in blue spheres on panels (A-C). Mutation L455S in JN.1 is shown in red spheres on panel A. The positions of KP.2 mutations R346T, L455S and F456L are shown in red spheres on panel B. The positions of KP.3 mutations L455S, F456L and Q493E are shown in red spheres on panel C.

The structural alignment of the predicted conformational ensembles (Figure 6) revealed only moderate heterogeneity of the flexible RBD loops while the RBD core remained largely rigid. Moreover, structural analysis of the AF2-predicted ensemble conformations with high pLDDT value indicated that the degree of mobility in the RBD loops, particularly the RBM binding motif, can remain fairly similar across all variants. However, we noticed that JN.1 which is the immediate descendant of the BA.2.86 variant displayed very small deviations from the BA.2.86 cryo-EM structure (Figure 6A), including RBD loop 444-452. This is also evident from mapping of the mutational positions in all predicted conformations, showing that V445H, G446S, Q493, L455S, F456 sites remain virtually in the same positions in the top predicted conformations (Figure 6A). The alignment of JN.1 RBD conformations revealed only moderate plasticity of the RBM motif which are reflected in minor lateral displacements of F486P and N481K positions (Figure 6A). We observed a generally similar pattern in the KP.2 RBD conformations where moderate variability was seen in V445H/G446S sites, but the predicted positions of N481K and F486P remain largely the same despite the overall mobility of the RBM region (Figure 6B). Interestingly, the RBD backbone conformations around L455S and F456L remains mainly rigid with only minor variations of the side-chains. Hence, functional heterogeneity of the RBD conformations may remain largely similar, suggesting that virus exploits convergent mutational sites to only moderately modulate ACE2 binding affinity, while the major role of these mutations is likely to enhance the immune evasion potential. A greater conformational heterogeneity of the functional RBD conformations was observed in the KP.3 RBD-ACE2 complex (Figure 6C). In this case, we found that the RBD loops 444-452 and 475-487 may display functionally relevant plasticity, where the RBB conformations and positions of F486P/N481K could slightly differ among top conformations (Figure 6C). Although the majority of mutational sites remain in similar positions in different conformations of the ensemble, we found some moderate plasticity associated with convergent mutations R346T, L455S, F456L and Q493E as well as in their interacting ACE2 sites K31, H34, E35. These findings from AF2 predictions suggest that L455S, F456L and Q493E in JN.1, KP.2 and KP.3 variants can undergo functional displacements around their native positions. Accordingly, it would indicate that respective changes in the binding affinities with ACE2 may be partly affected by concerted conformational rearrangements at the binding interface.

### 3.3 MD Simulations and Ensemble-Based Mutational Profiling of the Biding Interface Residues in the JN.1, KP.2 and KP.3 RBD-ACE2 Complexes

We performed comparative all-atom MD simulations of the BA.2.86, JN.1, KP.2 and KP.3 RBD-ACE2 complexes. The initial structure were taken from AF2 predicted ensembles. We analyze MD simulations using a comparative analysis of the residue-based distance fluctuation stability indexes (Supporting Information, Figure S2). The distributions showed that the local maxima for all RBD-ACE2 complexes are aligned with structurally stable and predominantly hydrophobic regions in the RBD core (residues 400-406, 418-421, 453-456) as well as key binding interface clusters (residues 495-505) that include binding hotspots R498 and Y501. The RBD positions associated with the high distance fluctuations stability indexes are F400, I402, Y421, Y453, L455, F456, Y473, A475, and Y489 (Supporting Information, Figure S2). The stability hotspots Y449, Y473, and Y489 are constrained by the requirements to maintain RBD stability and binding with the ACE2 host receptor. The profile minima are associated with the flexible RBD regions (residues 355-375, 381-394, 444-452, 455-471, 475-487). Despite a similar shape of the distributions for all Omicron RBD variants, the larger peaks were seen for JN.1 distribution (Supporting Information, Figure S2). This implies that both the RBD core and ACE2-binding interface positions are more rigid in JN.1 and become somewhat more flexinle in KP.2 and especialy KP.3 variants. The profiles pointed to flexible RBD regions (residues 355-375, 381-394, 444-452, 455-471, 475-487) that are shared by all variants and are associated with low stability indexes. Of particular interest are the lowered distance fluctuation stability indexes in the RBD regions 440-460 and 470-487 for KP.2 and KP.3 (Supporting Information, Figure S2). These observations are consistent with AF2 predictions suggesting a generlly more flexible RBD in the KP.2/KP.3 RBD variants. By mapping BA.2.86 mutational sites on the distriutions, we noticed that mutated RBD positions are typicaly characterized by moderate stability indexes, indicating that Omicron mutations target conformationally adaptable regions in the RBD. Mutational positions N440K, V445P, G446S, N460K, F486P displayed low distance fluctuation stability indexes, which may indicate presence of local mobility in these regions. Several important RBD binding interface centers L455 and F456 showed intermediate stability indexes, indicanting that although these positions remain overall stable in all variants, these sites may undero moderate fluctuations in their mutational forms containing L455S and F456L changes (Supporting Information, Figure S2). At the same time, R498, Y501 and H505 featured high stability indexes, reflecting a considerable rigidification of these residues due to strong interactions with ACE2.

We fist examined in some detail the predicted binding interfaces in the JN.1, KP.2 and KP.3 complexes. The binding interface in the cryo-EM structure of the BA.2.86 RBD-ACE2 complex (pdb id 8QSQ) showed that positions of H34 and K31 ACE2 residues as well as RBD residues Q493, L455 and F456 are very similar to those in the XBB.1.5 RBD-ACE2 complex [111] (Figure 7A, Supporting Information, Figure S3). In the XBB.1.5 RBD-ACE2 complex (pdb id 8WRL) K31 side chain is placed between Q493 and hydrophobic L455 and F456 residues, showing that movements of H34 and K31 are fairly restricted (Supporting Information, Figure S3A). A similar interfacial pattern was predicted for XBB.1.5 +L455F mutant [65,66] where the favorable position of Q493 remained largely unchanged as compared to XBB.1.5 (Supporting Information, Figure S3B). Compared to XBB.1.5, F456L mutation does not substantially affect the interactions on the RBD-ACE2 interface and yet Q493 and H34 undergo some rearrangements due to the increased conformational space which allows insertion of H34 side chain between Q493 and S494 [111] (Supporting Information, Figure S3C). A similar interfacial pattern is further strengthened in the XBB.1.5 +FLip RBD-ACE2 complex (pdb id 8WRH) [111] the flip of L455 and F456 provided synergistic reorganization of the interactions between H34 and Q493/S494 RBD residues (Supporting Information, Figure S3D).

**Figure 7.**
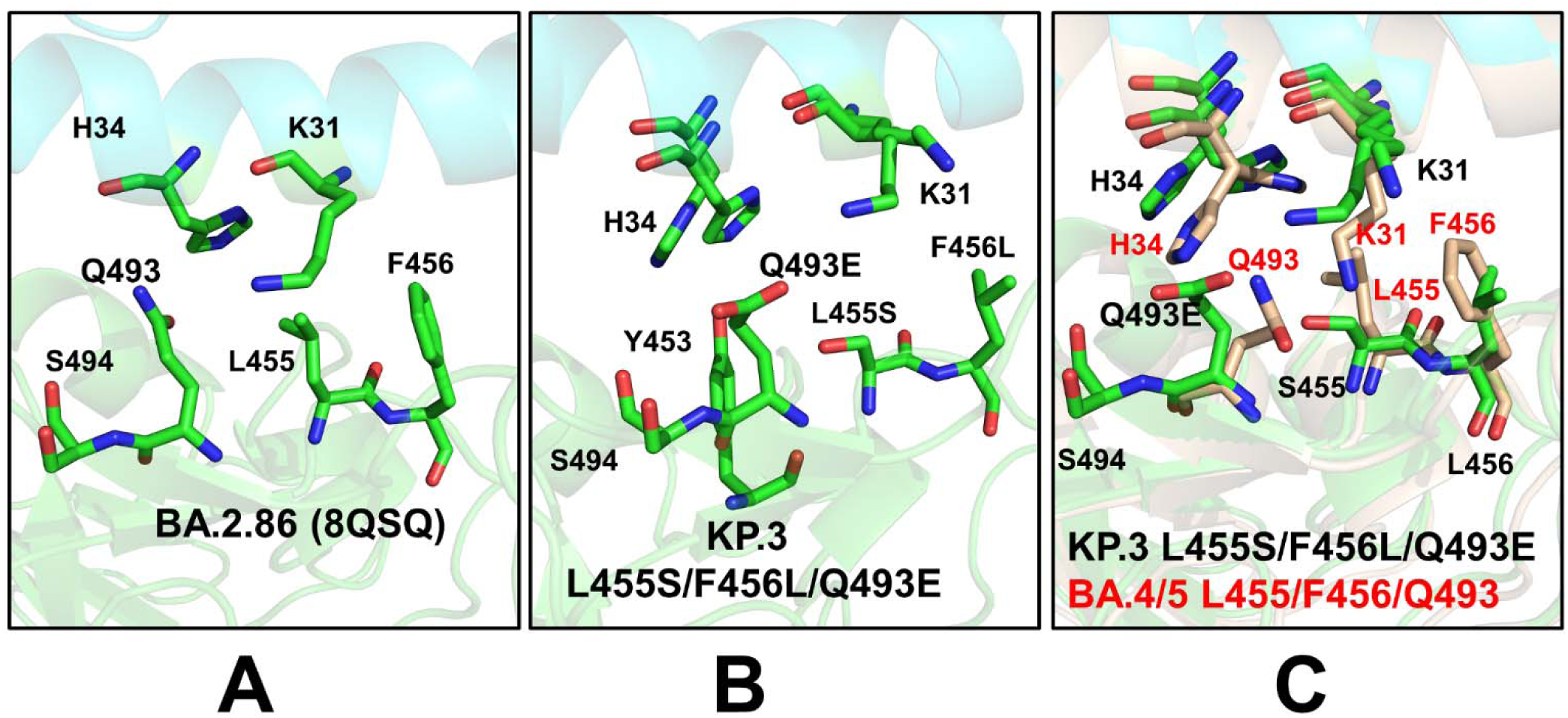
Structural predictions of the RBD-ACE2 binding interface. (A) A closeup of the AF2-predicted RBD binding interface residues Q493, S494, L455, F456, and ACE2 K31 and H34 for the BA.2.86 RBD-ACE2 complex. (B) A closeup of the AF2-predicted binding interface residues Q493E, S494, L455S, F456L and ACE2 K31, H34 for the KP.3 (L455S/F456L/Q493E) RBD-ACE2 complex. (C) A closeup of the binding interface residues Q493E, S494, L455S, F456L for KP.3 RBD-ACE2 complex overlayed on BA.4/BA.5 RBD-ACE2 complex The AF2-predicted binding interface residues are in atom-colored sticks. The binding interface residues S494, Q493, L455, F456, K31-ACE2 and H34-ACE2 are shown in brown sticks.

We examined the details of the predicted binding interface in the vicinity of L455 and F456 positions to illustrate the effect of mutations of L455 and F456 on the structural repacking induced by the L455F/F456L mutation and change in the binding contacts. Of special interest is the analysis of the binding interface in the KP.3 RBD-ACE2 complex, primarily to examine whether Q493E mutation can induce potential cooperative structural changes that may result in the observed epistatic improvement in the ACE2 binding affinity. Q493E involves the rarest of all nucleotide mutations, C->G, and occurs at a key residue. Intriguingly, Q493E marks another major change by reversing the basic trend in which the RBD interfacial region (438-506) has more basic i.e. positively charged in most of the Omicron variants. In fact, with V445H, N481K, and A484K mutations in BA.2.86 this variant exemplified largest increase in the positively charged RBD sites. Quite unexpectedly, Q493E, in the KP.3 variant reverses this trend, making the RBM more acidic. We leveraged the conformational ensembles obtained from AF2 and MD simulations to map the most favorable interfacial interactions in the KP.2 RBD-ACE2 complex (Figure 7B). The second region of the RBD that includes important mutational positions R403, L455, Y453, and Q493 binds with the central segment of the α1 helix interface of the ACE2 interface residues (D30, K31, H34, E35 and D38). Structural analysis showed that both H34 and K31 ACE2 residues become more flexible in the complex as compared to XBB.1.5 and BA.2.86 complexes (Figure 7B). Moreover, we found that H34 and K31 can assume several preferential side-chain orientations that enable moderate structural rearrangements which allow more room for making stable hydrogen bond interactions with Q493Q (Figure 7B). In addition, it is observed that the H34 side-chain placements allow for stronger contacts with Y453 residue. The additional comparative analysis illustrated superposition of the predicted binding interface for KP.3 RBD-ACE2 complex with the recently determined structures of the BA.4/5 RBD-ACE2 complex (pdb id 8H06) [112] (Figure 7C). Interestingly, the side chain of H34 adopts two alternative conformations in the BA.4/5 complex that are essentially identical to the preferential side-chain conformation of H34 in the KP.3 complex. H34 of hACE2 forms hydrogen bonding with Y453 of the BA.4/5 RBD and Q493 is hydrogen-bonded to K31 [112]. Similarly, the predicted K31 side-chain positions in the KP.3 complex are very similar to the experimentally determined K31 orientation in the BA.4/5 complex (Figure 7C).

Hence, structural analysis of the ensemble-predicted binding interface for the KP.3 complex suggested the moderately increased plasticity of the ACE2 interfacial sites that are afforded by L455S/F456L mutations that reduced bulkiness of the hydrophobic packing and conferred room for the increased flexibility and improved interactions mediated by Q493E with H34 and K31 hotspots (Figure 7C). In addition, we found that Q493 lies in the structural proximity of RBD residues 447-457, 467-473, 484-484, 488-497. In particular, Q493E is close to residues 346 (R346T in FLiRT variants), 448-456, and the 483-484 region where BA.2.86 has ΔV483 and A484K.

Using conformational ensembles obtained from MD simulation studies of the RBD-ACE2 complexes for the BA.2.86, JN.1, KP.2 and KP.3 variants we performed a systematic mutational scanning of the RBD residues in the RBD-ACE2 complexes. Mutational scanning of the key RBD positions in the cryo-EM structure of the BA.2.86 RBD-ACE2 complex (Supporting Information, Figure S4) was analyzed focusing on the RBD positions that correspond to sites of convergent evolution and determine changes seen in JN.1 (L455S), KP.2 (R346T, L455S, F456L) and KP.3 (L455S, F456L, Q93E). Most of mutations produced large destabilization binding free energy values with the exception of Y453F that resulted in the negligible change (Supporting Information, Figure S4A). This is in full agreement with the experimental studies that demonstrated role of Y453F as an adaptive mutation which increased virus interaction with mink ACE2 receptor, without compromising its utilization of human ACE2 receptor [113,114]. Mutational profiling at L455 showed that all modifications, particularly L455S are appreciably deleterious for ACE2 binding (ΔΔG ∼ 1.0 kcal/mol) (Supporting Information, Figure S4B). Mutational changes at F456 position of the BA.2.86 RBD-ACE2 structure are generally very unfavorable leading to ΔΔG ∼ 1.5-2.0 kcal/mol. However, F456L mutational change alone in the genetic background of BA.2.86 is rather small (ΔΔG ∼ 0.3 kcal/mol) and within the error margin, indicating that even individually F456L cannot significantly decrease the ACE2 binding affinity (Supporting Information, Figure S4C). Of particular interest are mutations in Q493 position showing relatively moderate destabilization for Q493R and Q493E (ΔΔG ∼ 0.6 kcal/mol) (Supporting Information, Figure S4D). These results agree with DMS studies that projected Q493E as destabilizing mutation in the backgrounds of XBB.1.5 variant. Our results suggested that F456L and Q493E may be only marginally deleterious and may reverse their effect on ACE2 binding.

We then leveraged AF2 and MD conformational ensembles to perform ensemble-based mutational scanning of key RBD positions that are mutated in the JN.1, KP.2 and KP.3 (Figure 8). Mutational analysis of S455 in the JN.1 RBD-ACE2 ensemble showed that mutations S455F, S455L and S455P are favorable and the reversed S455L change can potentially lead to some improvement in binding affinity (Figure 8A). This further confirmed that L455S mutation can markedly decrease ACE2 binding which is consistent with the experimental data [34,35]. We then investigated more closely the effect of mutations in key sites of KP.2 (R346T/L455S/F456L) KP.3 variant (S455, L456 and E493). The results showed that S455F, S455L and S455P remain favorable which implies that L455S substitution is deleterious for ACE2 binding in different genetic backgrounds of JN.1 and KP.3 (Figure 8B,D). Moreover, the L456 position in both KP.2 and KP.3 RBD-ACE2 complexes appears to be highly favorable as all substitutions lead to destabilizing ΔΔG ∼ 0.5-2.0 kcal/mol (Figure 8C,E).

**Figure 8.**
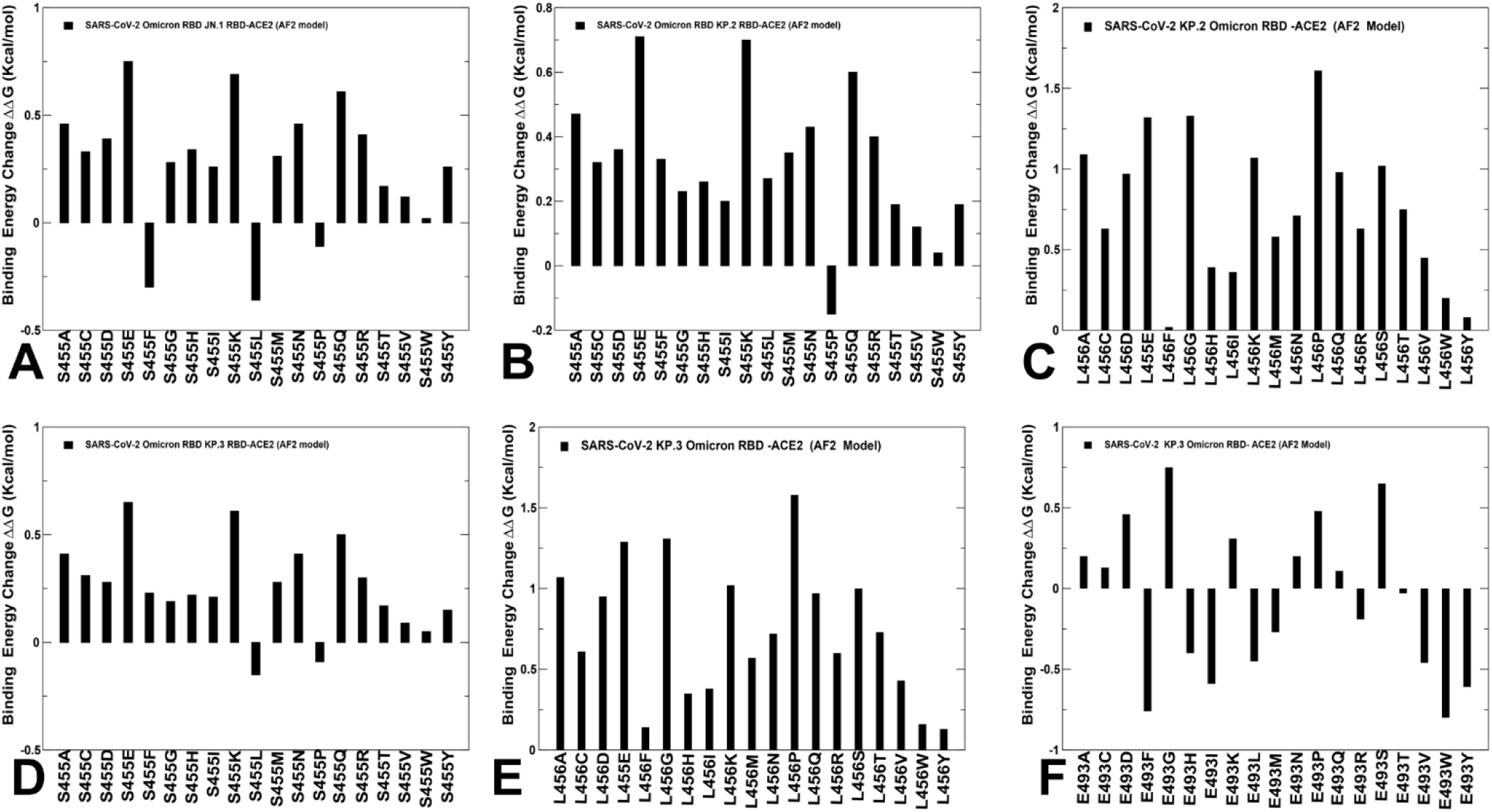
Ensemble-based mutational scanning of binding for the key RBD positions in backgrounds of JN.1, KP.2 and KP.3 variants. The conformational ensembles used in calculations are obtained from AF2 simulations and MD simulations of the RBD-ACE2 complexes for JN.1, KP.2 and KP.3 variants. The AF2 predicted structures were used for MD simulations. The profiles of computed binding free energy changes ΔΔG upon 19 single substitutions for S455 position in JN.1 (A), S455 in KP.2 (B), L456 in KP.2 (C), S455 in KP.3 (D), L456 in KP.3 (E) and E493 in KP.3 background (F). The respective binding free energy changes are computed using conformational ensembles obtained from AF2 predictions and MD simulations. The binding free energy changes are shown in black-colored filled bars. The positive binding free energy values ΔΔG correspond to destabilizing changes and negative binding free energy changes are associated with stabilizing changes.

Mutational profiling of E493 in the KP3 RBD-ACE2 complex showed a more complex and diverse picture as a number of modifications can improve binding affinity (Figure 8F) which is consistent with the fact that E493 is generally destabilizing change. However, in the KP.3 background, the reverse mutation E493Q appeared to be marginally destabilizing. i.e. E493 becomes slightly more favorable when combined together with KP.3 changes L455S and especially F456L (Figure 8F). Previous studies showed that salt bridge interactions in the BA.2 RBD formed by R493 with E37, E35 and D38 residues in ACE2 are partially lost in the BA.2.86 RBD-ACE2 complex [63]. In the BA.2.86 RBD-ACE2 complex Q493 forms stable interfacial contacts with D30, K31, N33, H34 and E35 including hydrogen-bonding interaction with the K31 side chain and the carboxyl group of E35 from ACE2 [63]. We showed that Q493E can partially rearrange the interactions with more flexible H34, E35 and K31 side-chains. Notably, the salt bridge formed by R493 of BA.2 with E35 of ACE2 is replaced by Q493 in BA.2.75, BF.7, XBB.1 and BA.2.86 [63]. Although mutational profiling of E493 in the KP.3 ensemble showed that many changes could be stabilizing, the observed changes are small and are often stabilizing for small hydrophobic substituents (Figure 8F). Overall, the results suggested that position 493 is quite tolerant to modifications and can be mutated in different Omicron variants to balance ACE2 binding and immune evasion. These results are also consistent with the experiments showing that small hydrophobic amino acids for S-Q493 and aromatic amino acids for S-N501 can be highly enriched for ACE2 binding, as these changes are expected to increase local hydrophobic packing [115]. This is also in agreement with the DMS experiments [43,44,47]. It was conjectured that the fact that these mutations predicted to increase ACE2 binding are not enriched in circulating SARS-CoV-2 variants suggests the affinity of the virus for its receptor is already sufficient for high transmission [115]. Our analysis also confirmed that R493/Q493/E493 in different Omicron variants including latest KP.3 can partially reassemble the interaction network while preserving convergent binding pattern of interactions which may enable evolutionary advantage through epistatic couplings between key hotspots at 456 and 493 positions on the RBD.

### 3.4. MM-GBSA Analysis of the Binding Affinities for the XBB RBD-ACE2 Complexes

According to the recent experimental studies using accurate SPR measurements the binding affinity of JN.1 RBD-ACE2 complex is K_D_ = 13 nM is reduced as compared to that of BA.2.86 variant (K_D_ = 1.7 nM) which is attributed to a deleterious L455S change in JN.1 [41]. At the same time, mutations F456L (K_D_ = 12 nM) and R346T + F456L (K_D_ = 11 nM) could not largely affect the ACE2-binding affinity of JN.1 indicating that the dampened ACE2 affinity of JN.1 due to L455S could not be fully compensated by F456L [41]. The central finding of this illuminating experimental study is that the Q493E mutation of KP.3 can markedly improve the ACE2 binding affinity revealing K_D_ = 6.9 nM which is only marginally lower than the superior binding affinity of the BA.2.86 variant [41]. Using the conformational equilibrium ensembles obtained fromAF2 predictions and MD simulations we computed the binding free energies for the RBD-ACE2 complexes using the MM-GBSA method [104–109]. To reduce noise and cancel errors in MM-GBSA computations we ran MD simulations on the complexes only, with snapshots taken from a single trajectory to calculate each free energy component. We computed and compared the RBD-ACE2 binding affinities for BA.2.86, JN.1 (BA.2.86+L455S), BA.2.86+F456L, BA.2.86+Q493E, BA.2.86+L455S, BA.2.86+F456L/Q493E, KP.2 (BA.2.86+L455S/F456L/R346T), and KP.3 (BA.2.86+L455S/F456L/Q493E) (Figure 9, Table 2). The results of MM-GBSA computations showed a robust agreement with the experimental binding affinities (Figure 9). The total binding free energy changes showed a more favorable binding affinity for the BA.2.86 RBD-ACE2 with the ΔG = −42.9 kcal/mol (Figure 9A).

**Figure 9.**
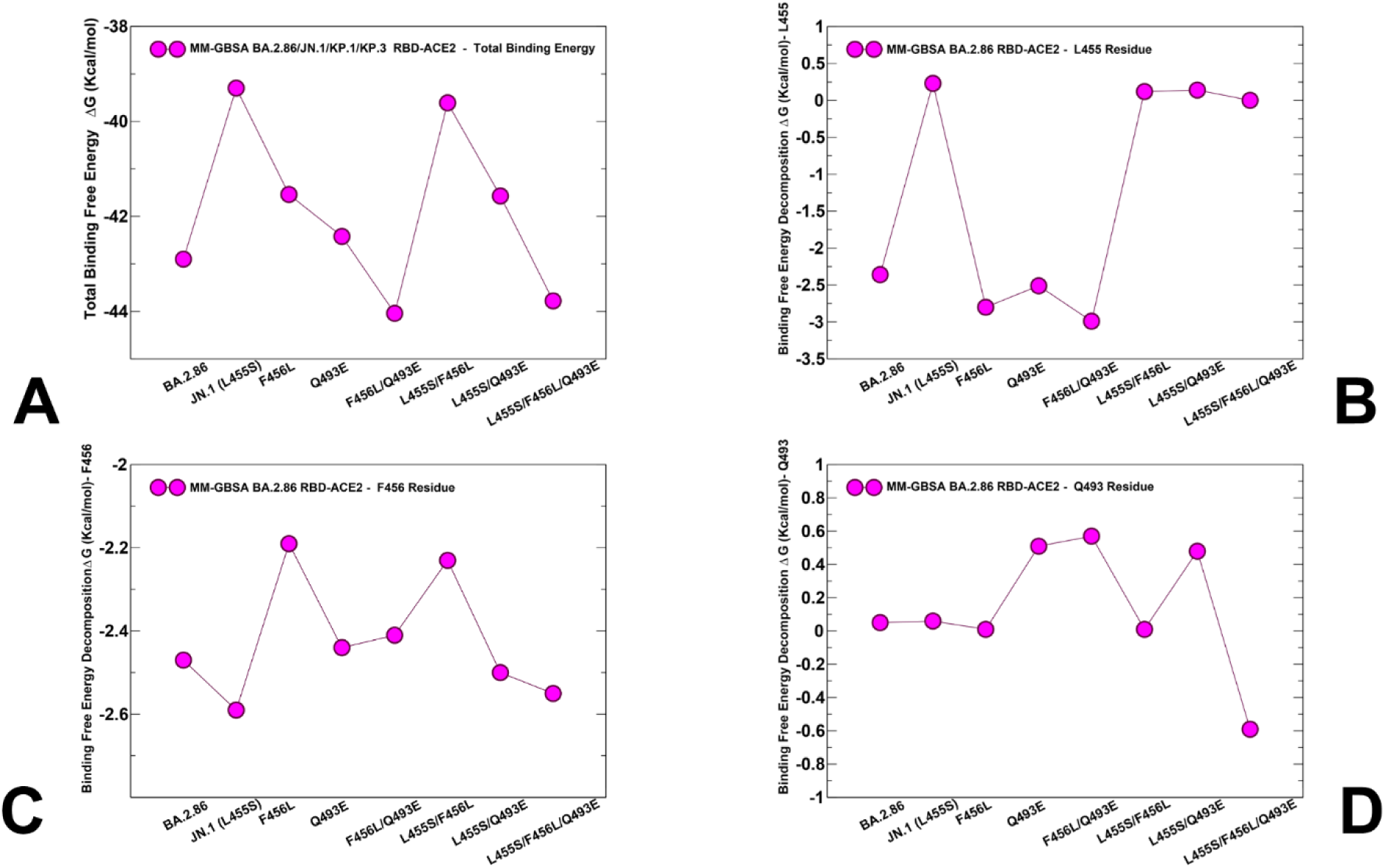
MM-GBSA binding free energies for BA.2.86, JN.1, BA.2.86+F456L, BA.2.86+Q493E, JN.1, BA.2.86+F456L/Q493E, KP.2, BA.2.86+L455S/Q493E and KP.3 RBD-ACE2 complexes. (A) The total binding free energies for the RBD-ACE2 complexes. (B) The MM-GBSA decomposition contribution of the binding energies for L455 position. (C) The MM-GBSA decomposition contribution of the binding energies for F456 position. (D) The MM-GBSA decomposition contribution of the binding energies for Q493E position.

**Table 2.** MM-GBSA Binding Energies for the BA.2.86, JN.1, KP.2 and KP.3 RBD-ACE2 Complexes. The experimetal binding contants are from [48])

| **System** | $E_{vdW}$ | $E_{elec}$ | $E_{GB}$ | $E_{SA}$ | $\Delta G_{bind}$<br>(kcal/mol) | Binding<br>constants<br>(nM) |
| --- | --- | --- | --- | --- | --- | --- |
| BA.2.86 RBD-ACE2 | -83.57 | -1438.64 | 1489.54 | -10.23 | -42.9 | 1.7 nM |
| BA.2.86+L455S (JN.1)<br>RBD-ACE2 | -79.2 | -1438.12 | 1487.67 | -9.66 | -39.3 | 13.0 nM |
| BA.2.86+F456L RBD-ACE2 | -80.91 | -1439.91 | 1489.21 | -9.93 | -41.54 | 12.0 nM |
| BA.2.86+Q493E RBD-ACE2 | -79.26 | -1222.65 | 1269.94 | -10.45 | -42.42 | unknown |
| BA.2.86+F456L/Q493E<br>RBD-ACE2 | -79.49 | -1224.61 | 1270.23 | -10.17 | -44.04 | unknown |
| BA.2.86+L455S/F456L<br>(KP.2) RBD-ACE2 | -77.40 | -1440.2 | 1487.22 | -9.31 | -39.61 | 11.0 nM |
| BA.2.86+L455S/Q493E<br>RBD-ACE2 | -77.2 | -1222.66 | 1268.2 | -9.97 | -41.57 | 21.0 nM |
| BA.2.86+L455S/F456L/Q493E (KP.3) RBD-ACE2 | -77.96 | -1226.20 | 1269.98 | -9.60 | -43.78 | 6.9 nM |

Consistent with the experiments, we found that L455S in JN.1 can lead to significant reduction in binding with ACE2 as compared to BA.2.86, showing ΔG = −39.3 kcal/mol (Figure 9A). As may be expected, the breakdown of the MM-GBSA binding energies (Table 2) showed the key role of the favorable van der Waals interactions (ΔG = −83.57 kcal/mol) in BA.2.86 that is markedly more favorable than that in JN.1 (ΔG = −79.20 kcal/mol). This MM-GBSA analysis confirmed that L455S in JN.1 can reduce the interaction energy and reduce the binding affinity. The energetic analysis of the BA.2.86+F456L complex revealed that F456L alone has only minor effect on the binding affinity resulting in the overall ΔG = −41.54 kcal/mol (Figure 9A, Table 2) which is comparable to that of the BA.2.86 RBD-ACE2 complex. These results agree with reported unpublished observations by Starr (https://x.com/tylernstarr/status/1800315116929560965).

According to MM-GBSA calculations, the binding affinity of the BA.2.86+Q493E and especially BA.2.86+Q493E/F456L complexes displayed a revealing and interesting trend in which the binding free energy is progressively improved to ΔG = −42.42 kcal/mol and ΔG = −44.04 kcal/mol (Figure 9, Table 2). Hence, our predictions indicate that coupling of Q493E and F456L mutations may fully restore binding affinity of the BA.2.86 variant. Interestingly, the MM-GBSA breakdown of the energetic components showed the reduced electrostatic contribution in variants bearing the Q493E mutation, but this reduction is compensated by the decreased polar solvation penalty, leading to the overall more favorable binding energies. The comprehensive MM-GBSA analysis is also in line with the experiments [42] by demonstrating that binding affinity of the BA.2.86 L455S/F456L variant (ΔG = −39.61 kcal/mol) is similar to that of JN.1 (BA.2.86+L455S) and therefore suggesting that F456L mutation alone cannot compensate the loss of binding due to L455S. The important finding of our analysis is that KP.3 variant that bears L455S, F456L and Q493E displayed a considerably improved binding affinity (ΔG = −43.78 kcal/mol) as compared to the individual effects of F456L and Q493E alone (Figure 9A, Table 2). These results are consistent with the very latest experimental data showing that non-additive epistatic interactions between Q493E and other mutations of KP.3 can represent the energetic driver behind unexpectedly enhanced affinity of KP.3 [41].

To evaluate the contributions of L455, F456 and Q493 residues and corresponding mutations across all studied variants, we also reported the MM-GBSA residue-based breakdown (Figure 9B,C,D). We found that L455S can induce consistent and similar loss of binding interactions across all variants that share this mutation (Figure 9B), thus indicating that deleterious effect of L455S is not affected by presence of F456L and Q493E mutations. Of particular interest is MM-GBSA decomposition analysis at 456 position (Figure 9). We noticed that F456L can induce loss of binding interactions in the BA.2.86+F456L, while partly reversing the trend and restoring more favorable contribution in BA.2.86+F456L/Q493E and KP.3 (BA.2.86+L455S/F456L/Q493E) variants (Figure 9C). An overall similar trend is seen in the MM-GBSA decomposition of Q493 position (Figure 9D) where small improvements in the contribution of 493 position could be observed in the KP.3 variant. However, the differences between contributions of Q493/E493 in different variants are relatively minor. We also evaluated contributions of individual ACE2 positions H34 and K31 that are involved in interactions with L455, F456 and Q493 sites (Supporting Information, Figure S5). It can be seen that that BA.2.86 variants with F456L/Q493E and L455S/F456L/Q493E additions lead to stronger contribution of K31 position on ACE2 due to favorable combination of van der Waals and electrostatic interactions (Supporting Information, Figure S4A). In BA.2.86 Q493 is hydrogen-bonded with K31 while in F456L/Q493E and L455S/F456L/Q493E variants, E493 and K31 form strong ionic electrostatic interactions and there are more van der Waals contacts between Q493E and K31 of ACE2 (Supporting Information, Figure S5A). We also found that contribution of H34 becomes more favorable in variants sharing F456L ((Supporting Information, Figure S5B). Moreover, multiple side-chain conformations of H34 are involved in multiple contacts and form favorable interactions with Y453.

Together, structural and energetic analysis of BA.2.86 combinations including JN.1, KP.2 and KP.3 variants suggested that F456L and Q493E may act cooperatively in KP.3 to induce epistatic couplings and restore the binding affinity to the level of BA.2.86. Although precise molecular mechanisms underlying the observed epistatic effects of Q493E and F456L are likely to be complex, our results suggested that the increased flexibility of the RBD in the KP.3 complex may allow for more plasticity of the reciprocal ACE2 sites H34 and K31 that appeared to adopt multiple side-chain conformations in the structural ensembles. We found the side chain of H34 adopts two alternative conformations that are also seen in the BA.4/5 complex and forms hydrogen bonding with Y453 [112]. Structural examination of the predicted ensembles and MM-GBSA analysis of the RBD-ACE2 complexes for JN.1, KP.2 and KP.3 variants suggested that the plasticity of the ACE2 interfacial sites that are afforded by L455S/F456L mutations can be properly exploited by Q493E, Y453 and F456L positions to enhance binding affinity. Overall, this analysis pointed to a central role of F456L as a potential regulator of epistatic couplings but also suggests a potential role of structurally proximal L452 and Y453 sites supporting the improved binding interfacial interactions in the KP.3 variant.

### 3.5. Mutational Profiling of Protein Binding Interfaces with Distinct Classes of Antibodies

We embarked on structure-based mutational analysis of the S protein binding with different classes of RBD-targeted antibodies, focusing specifically on the role of BA.2.86 mutations in mediating potential resistance to broad class of antibodies and eliciting robust immune escape. We specifically examined a panel of monoclonal antibodies that were reported to retain activity against BA.2. XBB.1.5 and BA.2.86 variant but showed reduced neutralization against JN.1, KP.2 and KP.3 variants [41]. Structure-based mutational scanning of the S protein binding interfaces with a panel of class I RBD-targeting antibodies included S2K146 [71], Omi-3 [72] Omi-18[72], Omi-42 [72] as well as antibodies BD55-5514 (SA55) and BD55-5840 (SA58) [73,74] that bind to a different RBD epitope and are experimentally known to tolerate escape mutations in BA.2.8, JN.1, KP.2 and KP.3 variants. The results were compared with biochemical measurements from a panel of broadly neutralizing monoclonal antibodies [34,40,41]. In particular, the experiments found that L455S of JN.1 significantly enhances immune evasion at the expense of reduced ACE2 binding affinity [34].

To provide a systematic comparison, we constructed mutational heatmaps for the RBD interface residues of the S complexes with S2K146 (Figure 10A), OMI-3 (Figure 10B), OMI-18 (Figure 10C) and OMI-42 antibodies (Figure 10D). Strikingly, in all S complexes with class I antibodies mutational heatmaps clearly showed that a pair of adjacent residues L455 and F456 corresponding to convergent evolutionary hotspots are also dominant escape hotpots of antibody neutralization. These results are consistent with the experimental data showing that antibody evasion drives the convergent evolution of L455F/S and F456L, while the epistatic shift caused by F456L can facilitate the subsequent convergence of L455 and Q493 changes to restore ACE2 binding [41]. Furthermore, mutational heatmaps indicated that other important positions involved in JN.1, KP.2 and KP.3 variants, particularly Q493 site can be considered as a secondary escape hotspot sine mutations of Q493 across all class I antibodies (Figure 10). Interestingly, most of the RBD residues that directly bind S2K146 are also involved in binding to ACE2. Indeed, the mutational heatmap of the RBD binding against S2K146 identified that all substitutions in key interfacial positions can incur a consistent and considerable loss in binding affinity with S2K146 (Figure 10A).

**Figure 10.**
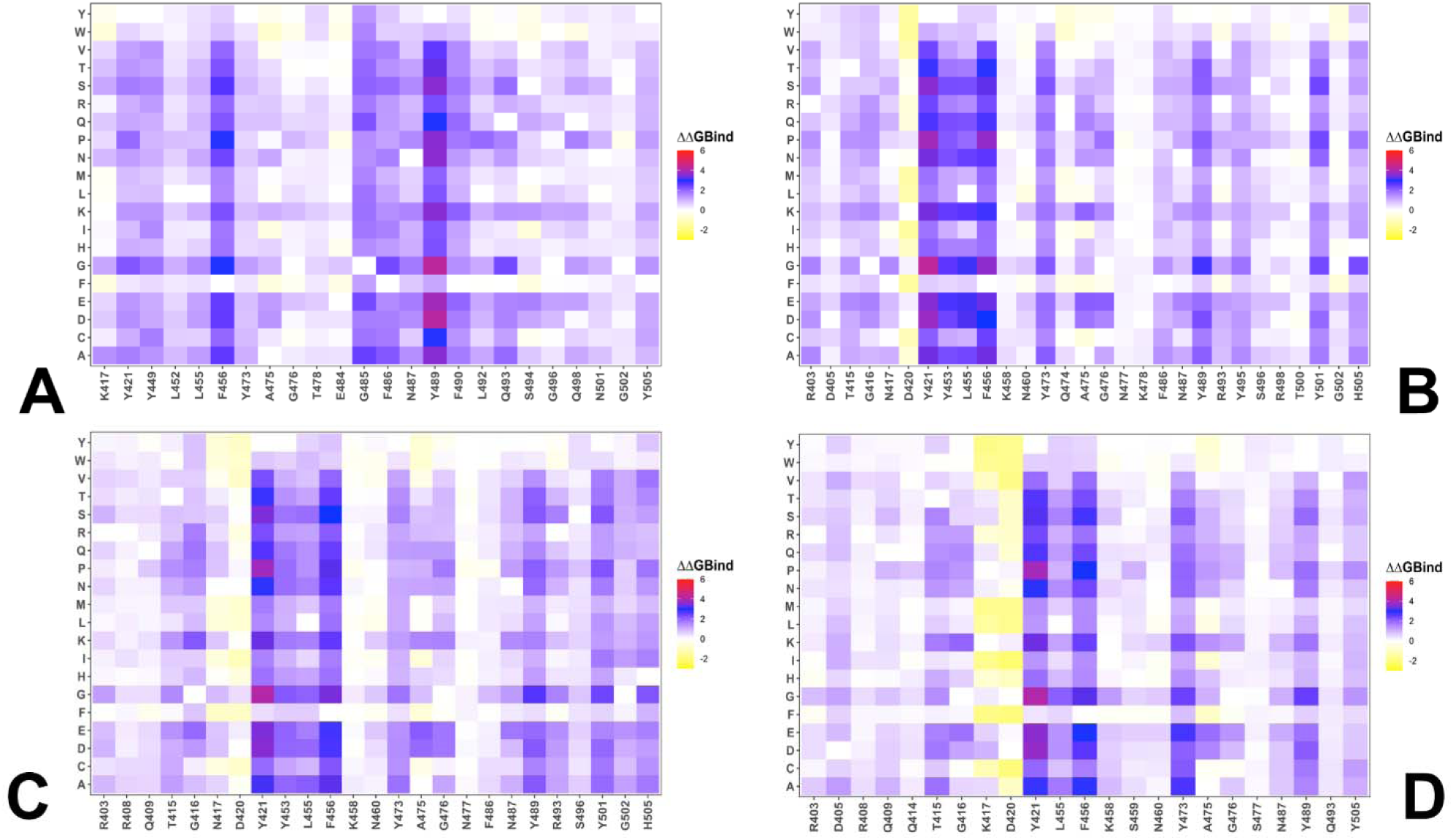
Ensemble-based dynamic mutational profiling of the RBD intermolecular interfaces in the Omicron RBD-ACE2 complexes. The mutational scanning heatmaps are shown for the interfacial RBD residues in the BA.2.86 RBD-ACE2 (A), JN.1 RBD-ACE2 (B), KP.2 RBD-ACE2 (C), and KP.3 RBD-ACE2 complexes (D). The heatmaps show the computed binding free energy changes for 20 single mutations of the interfacial positions. The standard errors of the mean for binding free energy changes using randomly selected 1,000 conformational samples (0.11-0.16 kcal/mol) were reduced to 0.08-0.12 kcal/mol when using equally distributed 10,000 samples from the MD trajectories.

Among these sites are F456, F486, N487, Y489, F490, and Q493 as mutations in these sites caused the largest losses in the binding affinity with S2K146 (Figure 10A). These findings are full consistent structural studies showing that the S2K146 footprint on the SARS-CoV-2 RBD mimics that of the ACE2 receptor, with 18 of 24 epitope residues shared with the ACE2 binding site, including L455, F486, Q493, Q498, and N501 [71]. The important finding of the mutational heatmap analysis is that positions Y421, Y453, L455 and F456 emerged as key escape hotspots in S binding with Omi-3, Omi-18 and Omi-42 class I antibodies (Figure 10B-D). L455 and F456 positions are located at the epitope of RBD Class 1 antibodies and neutralization assays demonstrated that L455S mutation enables JN.1 to evade Class 1 antibodies [34]. Mutational heatmaps data are consistent with these experiments, showing that effectively all modifications in L455, including L455S, can cause considerable loss in antibody binding. In addition, the results also highlighted the importance of Y453 position that is involved in favorable interactions with the ACE2 and antibodies. A secondary group of escape hotspots for these class I antibodies included F486, N487, Y489, and Q493 positions (Figure 10B-D). Mutations in some of these sites such as F486P are implicated as the main immune escape hotspots for BA.2.86 variant. Finally, due to mimicry of the ACE2 binding, other escape positions correspond to the ACE2 binding affinity hotspots Y501 and H505 (Figure 10). Importantly, the result firmly suggested that major drivers of immune escape for this class I antibodies correspond to L455 and F456 sites that undergo mutations in the JN.1, KP.2 and KP.3 variants, enabling evolution through the enhanced immune escape.

We then examined JN.1, KP.2 and KP.3 mutations using BA.286 as genetic background to quantify binding free energy changes of S binding with the class I antibodies (Figure 11). The binding free energy changes associated with BA.2.86 mutations in the complex with S2K146 (Figure 11A) showed an appreciable loss of binding upon K417N, L455, F456L, F486P, Q493E and Q498R mutations. Interestingly, the largest destabilization changes were induced L455S (ΔΔG = 1.03 kcal/mol), F456L (ΔΔG = 1.55 kcal/mol) and Q493E (ΔΔG = 1.59kcal/mol). Hence, these key mutational changes present in JN.1, KP.2 and KP.3 variants may induce progressively enhanced immune escape from S2K146 which is consistent with the experiments [34,40,41]. We found that L455S and F456L mutations induce very significant losses in antibody binding with Omi-3, Omi-18 and Omi-42 antibodies (Figure 11B-D). In particular, for binding with Omi-3, L455S mutation caused ΔΔG = 2.3 kcal/mol and F456L incurred ΔΔG = 1.76 kcal/mol (Figure 11B), while for Omi-42 these losses were ΔΔG = 1.62 kcal/mol for L455S and ΔΔG = 1.53 kcal/mol for F456 (Figure 11D). These results are consistent with the experiments showing the increased evasion against JN.1 variant with Omi-3 and Omi-18 antibodies [34,40,41] as L455 and F456 sites emerge as dominant hotspots where mutations L455S and F456L caused the largest loss in affinity. Common to both S2K146 and Omi-3 antibodies, we also observed a considerable loss of binding due to F486P mutation (Figure 11A,B). Specific for Omi-3 antibody is a destabilizing role of N460K mutation inducing loss of binding ΔΔG = 0.83 kcal/mol (Figure 11B). Interestingly, Q493E emerges as the third most important escape mutation in binding with Omi-3 and Omi-18 antibodies (Figure 11B,C). Structural mapping of the binding epitope residues and sites of BA.2.86 mutations (Figure 11E-H) highlighted similar binding modes for these antibodies, targeting the second region of the RBD that includes R403, L455, Y453, and Q493 sites. In general, the results confirmed the experimental finding that KP.2 and KP.3 harboring the F456L mutation, alone or in combination with Q493E (KP.3) can significantly impair the neutralizing activity of RBD class-1 monoclonal antibodies such as BD-1854, BD57-1302, and Omi-42 [40].

**Figure 11.**
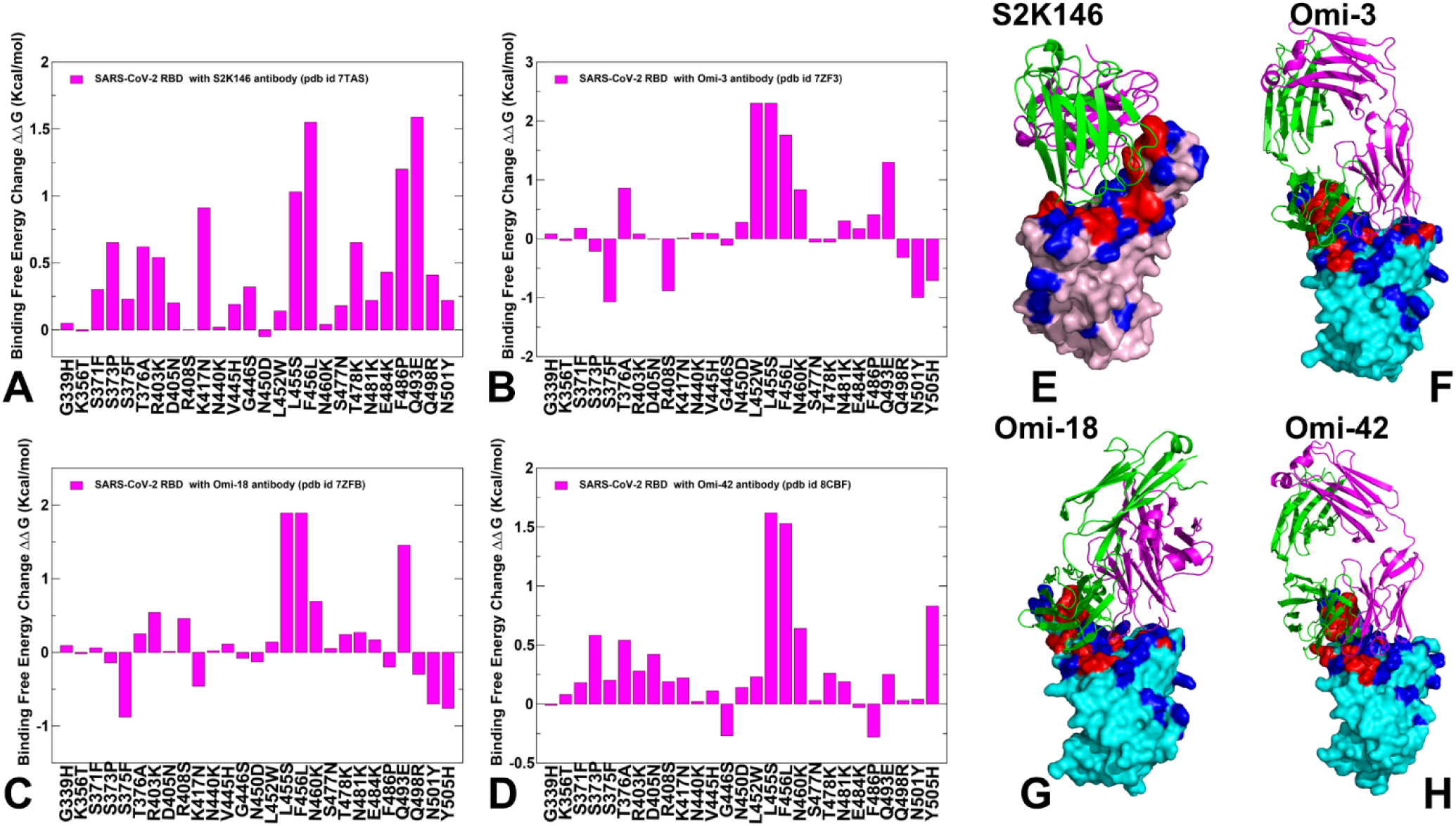
Structure-based mutational profiling of the S-RBD complexes with class 1 of RBD antibodies. The mutational screening evaluates binding energy changes induced by BA.2.86/JN.1/KP.2/KP.3 mutations in the RBD-antibody complexes. Mutational profiling of the S-RBD complex with S2K146, pdb id 7TAS (A), S-RBD Omicron complex with Omi-3, pdb id 7ZF3 (B), S-RBD Omicron complex with Omi-18, pdb id 7ZF8 (C), and S-RBD in complex with Omi-42, pdb id 8CBF (D). The binding free energy changes are shown in magenta-colored filled bars. The positive binding free energy values ΔΔG correspond to destabilizing changes and negative binding free energy changes are associated with stabilizing changes. The 3D structures of the RBD-antibody complexes are shown for RBD-S2K146 (E), RBD-Omi-3 (F), RBD-Omi-18 (G), and RBD-BD-515 complexes (H). The RBD is shown in cyan-colored surface representation. The RBD binding epitope residues are shown in red and BA.2.86 mutational positions are shown in blue. The antibodies are shown in ribbons (heavy chain in magenta and light chain in green-colored ribbons).

BD55-5514 belongs to class F2 and F3 antibodies that compete with ACE2, and their binding is affected by T376, K378, D405, R408 and G504, [73,74]. BD55-5514 displayed high potency against the Omicron subvariants and recently developed non-competing antibody cocktail of BD55-5840 (also known as SA58; class 3) and BD55-5514 (also known as SA55; class 1/4), displayed high potency against the Omicron subvariants [116]. Recent studies by Cao group showed that BD55-5514 (SA55) antibody can retain neutralizing efficacy against most of the known Omicron variants including JN.1 [34]. SA55 antibody also showed significant neutralization activity against other variants sharing F456L mutation including HV.1 (L452R+F456L), JD.1.1(L455F/F456L+A475V) that typically greatly increase antibody evasion for other class I antibodies at the cost of ACE2 binding. The detailed mutational heatmap of the BD55-5514 interactions with the S protein showed a very different picture as compared to class I antibodies such as Omi-3, Omi-18 and Omi-42 (Figure 12A). The map showed that the S373P and S375F mutations could promote the interaction with BD55-5514. T376, D405, and R408 are involved in the interaction with BD55-5514 but they are all located at the periphery of the BD55-5514 epitope, and mutations in these positions have moderate effect on binding affinity (Figure 12A). Importantly, BD55-5514 binding interface does not involve escaping hotspot positions of BA.2.86, JN.1 and KP.2/KP.3 such as L455, F456 and Q493 (Figure 12A). A more detailed profiling of JN.1/KP.3 mutations against BD55-5514 antibody showed only small destabilization changes upon mutations T376A (ΔΔG = 0.81 kcal/mol), R403K (ΔΔG = 0.65 kcal/mol), D405N (ΔΔG = 0.79 kcal/mol), R408S (ΔΔG = 0.34 kcal/mol), L455S (ΔΔG = 0.7 kcal/mol) and F456L (ΔΔG = 0.51 kcal/mol) (Figure 12B). These changes reflect mostly a mild loss in the RBD stability and binding interactions, which is consistent with functional experiments showing group F3 antibodies, such as BD55-5514, are not sensitive to the D405N and R408S mutations of BA.2 making this antibody effective against a broad spectrum of recent variants from BA.2.86 to KP.2 and KP.3 [116]. Structural mapping of the binding epitopes and BA.2.86 mutational sites for BD55-5514 and BD55-5840 antibodies (Figure 12 C,D) illustrated how BD55-5514 can bind to the RBD together with BD55-5840 without interference.

**Figure 12.**
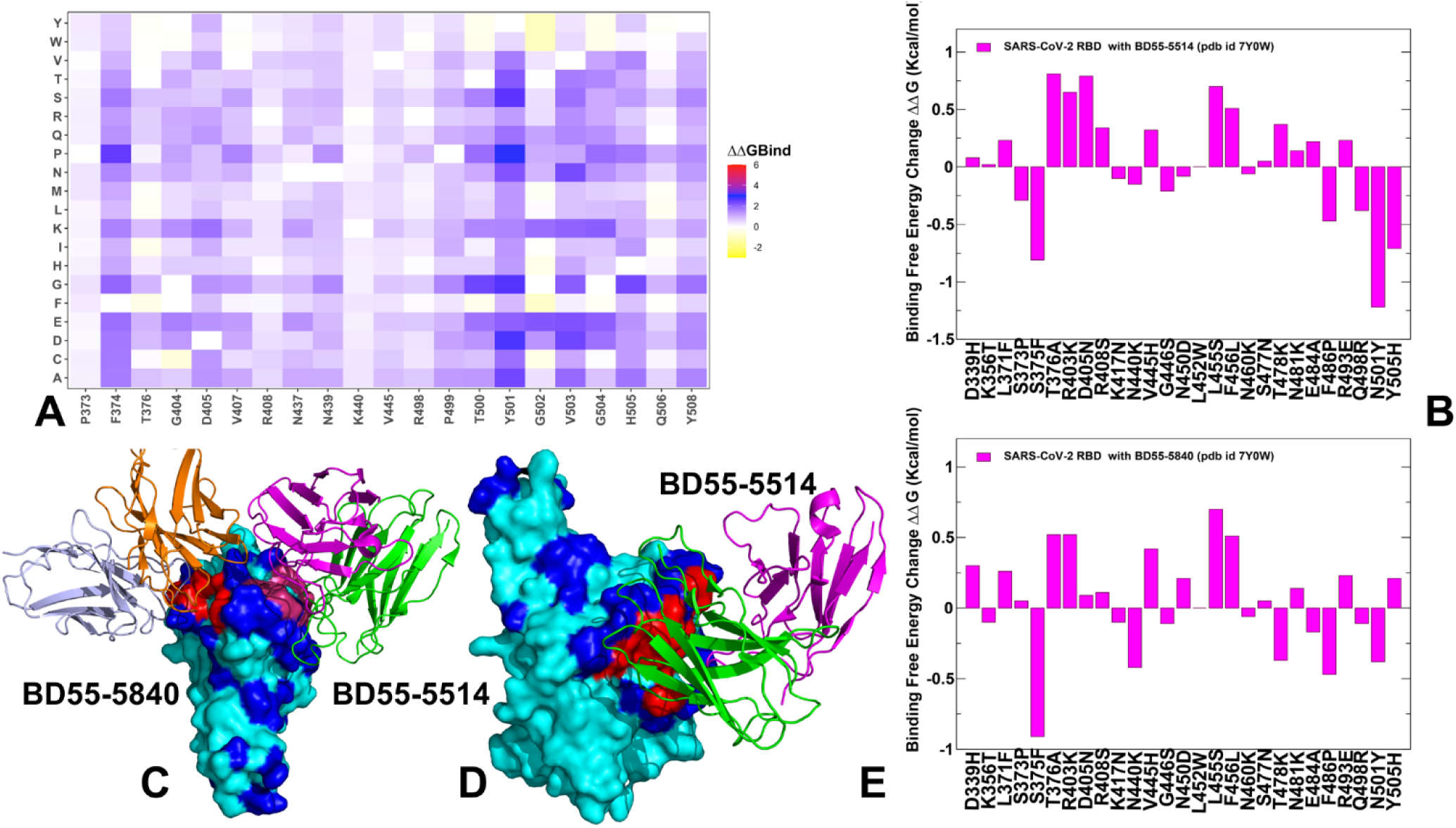
Structure-based mutational profiling of the S complexes with BD55-5840 (SA58; class 3) and BD55-5514 (SA55; class 1/4) antibodies. The mutational screening evaluates binding energy changes induced by BA.2.86/JN.1/KP.2/KP.3 mutations in the RBD-antibody complexes. (A) The mutational heatmap of the BD55-5514 interactions with the S protein shows the computed binding free energy changes for 20 single mutations of the interfacial positions. (B) Mutational profiling of the S complex with BD55-5514, pdb id 7Y0W. (C). The structure of the S complex with BD55-5840 and BD55-5514. (D) The structural view of the S protein bound to BD55-5514 in the same complex. The RBD is shown in cyan-colored surface representation. The RBD binding epitope residues are shown in red and BA.2.86 mutational positions are shown in blue. The antibodies are shown in ribbons (heavy chain in magenta and light chain in green-colored ribbons). (E) Mutational profiling of the S complex with BD55-5840, pdb id 7Y0W. The binding free energy changes are shown in magenta-colored filled bars. The positive binding free energy values ΔΔG correspond to destabilizing changes and negative binding free energy changes are associated with stabilizing changes.

Mutational analysis of the BD55-5514 and BD55-5840 interactions confirmed that S373P and S375F mutations can improve the binding affinities of these antibodies with the S protein which is also consistent with the pseudo-virus data [74]. We also found that binding of BD55-5840 antibody is even less affected by mutations T376A, R403K and D405N as compared to BD55-5514 (Figure 12E). Importantly, both L455S and F456L mutations are only marginally deficient for binding of these antibodies with ΔΔG ∼0.5-0.7 kcal/mol (Figure 12A,E). As a results, these antibodies could potently bind to the entire spectrum of BA.2.86, JN.1, KP.2 and KP.3 mutations. Our data enabled the quantitative energetic analysis of the binding interactions for these antibodies in the complex where BD55-5840 and BD55-5514 act synergistically and bind to different sides of the RBD (Figure 12C). Interestingly, structural mapping of the BA.2.86 mutational sites onto the RBD complex with BD55-5514/BD55-5840 highlighted that binding modes of antibodies do not significantly overlap with the mutational positions (Figure 12 C) and, as a result, this antibody combination can exhibit exceptional neutralizing activity by successfully avoiding immune escape hotspots. Together, structural and energetic analysis provides a rationale to the experimental results showing that BD55-5840 (SA58) + BD55-5514 (SA55) cocktail exhibits remarkable neutralization breadth and potency [74,116] and particularly SA55 can retain neutralizing efficacy against all examined variants BA.2.86, JN.1, KP.2 and KP.3 [34].

A very recent study introduced CYFN1006-1 and CYFN1006-2 antibodies that demonstrated consistent neutralization of all tested SARS-CoV-2 variants comparable to or even superior to those of SA55 [117]. CYFN1006-2 exhibited high potency against all SARS-CoV-2 variants, with a slightly reduced efficacy against KP.2 [117]. These antibodies have binding epitopes that overlap with LY-CoV1404, REGN10987 and S309 that are situated on the outer surface of RBD and bind to a different RBD region compared to SA55. Hence, a cocktail with combinations of SA55 and CYFN1006-1 may be potentially beneficial against JN.1, KP.2, KP.3 and evolving mutants of SARS-CoV-2 [117].

## 4. Discussion

The results of this study provided molecular rationale and support to the experimental evidence that functionally balanced substitutions that optimize tradeoffs between immune evasion, high ACE2 affinity and sufficient conformational adaptability might be a common strategy of the virus evolution and serve as a primary driving force behind the emergence of new Omicron subvariants, including but not limited to BA.2.86, JN.1, KP.2 and KP.3 variants. Recent studies of emerging Omicron variants, particularly evolving BA.2.86 sublineage suggested that the evolutionary paths for significant improvements in the binding affinity of the Omicron RBD variants with ACE2 are relatively narrow and may involve convergent mutational hotspots to primarily optimize immune escape while retaining sufficient ACE2 affinity. These mechanisms based on convergent adaptation may determine the scope of “evolutionary opportunities” for the virus to adapt new mutations that improve immune resistance without compromising ACE2 binding affinity and stability. Our multifaceted study explored quantitative aspects of structure, dynamics and energetics of the BA.2.86, JN.1 and KP.2/KP.3 variants, showing a robust agreement with a large body of experimental data on ACE2 binding and interactions with various classes of antibodies. The results of our investigation suggested that the existence of epistatic interactions between convergent mutational sites at L455, F456, Q493 positions that enable to protect and restore ACE2 binding affinity while conferring beneficial immune escape. Consistent with the latest functional studies, our results showed that Q493E and F456L can act cooperatively through epistatic couplings to reverse the detrimental effect of individual Q493E mutation seen in other genetic backgrounds. Although precise molecular mechanisms underlying the observed epistatic effects of Q493E and F456L are likely to be complex, our results suggested that epistatic interactions between these sites may arise due to the increased side-chain flexibility of the interacting F456L, Q493E on RBD with H34 an K31 on ACE2 within a rather confined RBD-ACE2 interface. The progressively increased flexibility of the RBD-ACE2 interface in the KP.2 and KP.3 complexes is manifested in multiple side-chain conformations for H34 and K31 seen in the structural ensembles as well as the enhanced plasticity of the binding interface afforded by L455S and F456L mutations. Overall, this analysis pointed to a central role of F456L as a potential regulator of epistatic couplings but also suggests a potential role of structurally proximal Y453 site supporting the improved binding interfacial interactions in the KP.3 variant. Our results are consistent with the related study showing that Omicron variants can induce distinct dynamics and exploit epistatic interactions between sites R346, F486P, Q498, Q493 to modulate the RBD-ACE2 interface with increasing flexibility in antibody binding residues [118]. The results also demonstrated that L455, F456 and Q493 are pronounced escape hotspots of resistance to class I antibodies. We suggested that epistatic couplings between these sites may not only help to restore ACE2 binding affinity but also represent a mechanism for amplifying immune response provided by individual mutations L455S, F456L and Q493E. These arguments are also consistent with evolutionary studies revealing strong epistasis between pre-existing substitutions in Omicron variants and antibody resistance mutations acquired during selection experiments, suggesting that epistasis can also lower the genetic barrier for antibody escape [119]. Based on the correspondence between the computational results and biochemical experiments we suggest that the primary role of BA.2.86, JN.1, KP.2, and KP.3 mutations may be to ensure a broad resistance against different classes of RBD antibodies, while several important mutations such as Q493E could confer the improved ACE2 binding affinity. The results of our study provide an additional evidence to the mechanism proposed by Bloom and colleagues that recently evolving variants tend to exploit a fairly focused group of antibody-escape sites (L455, F456, R403, D405) through convergent evolution as these positions are under moderate functional constraint and mutations incur little cost to virus. Although the effect of immune evasion could be more variant-dependent and modulated through recruitment of mutational sites in various adaptable RBD regions, the currently dominating variants operate on a limited number of convergent escape hotspots. At the same time, the epistatic interactions between convergent mutational sites and ACE2 binding affinity hotspots may represent a viable mechanism to retain ACE2 binding with minimum resources and leveraging the immune escape centers to protect ACE2 affinity.

## 5. Conclusions

In this study, we combined AF2-based atomistic predictions of structures and conformational ensembles of the SARS-CoV-2 Spike complexes with the host receptor ACE2 for the most recent dominant Omicron variants JN.1, KP.1, KP.2 and KP.3 to examine the mechanisms underlying the role of convergent evolution hotspots in balancing ACE2 binding and antibody evasion. The AF2-predicted conformational ensembles suggested the increased heterogeneity in the JN.1, KP.2 and especially KP.3 RBD variants which may potentially enable these variants to leverage a more mobile RBD structure to modulate and evade antibody neutralization. The results showed that the majority of mutational sites remain in similar positions in different conformations of the AF2 ensembles, but convergent mutational sites R346T, L455S, F456L and Q493E as well as in their interacting ACE2 sites K31, H34, E35 may exhibit some moderate level of plasticity and display concerted conformational rearrangements at the binding interface. Using conformational ensembles obtained from MD simulation studies of the RBD-ACE2 complexes for the BA.2.86, JN.1, KP.2 and KP.3 variants we performed a systematic mutational scanning of the RBD residues in the RBD-ACE2 complexes. Our results suggested that F456L and Q493E may be only marginally deleterious and may reverse their effect on ACE2 binding in a different genetic background. Mutational profiling in the KP.3 background showed that Q493E mutation becomes more favorable when combined together with KP.3 changes L455S and especially F456L. The results of our study supported a recently proposed hypothesis that KP.2 and KP.3 lineage may have evolved to outcompete other Omicron subvariants by improving immune suppression while balancing binding affinity with ACE2 via through compensatory epistatic effect of L455S, F456L, Q493E and F486P mutations. Based on structural analysis of the conformational ensembles and MM-GBSA computations of the RBD-ACE2 complexes for JN.1, KP.2 and KP.3 variants, we argue that the plasticity of the ACE2 interfacial sites that are afforded by L455S/F456L mutations can be properly exploited by Q493E, Y453 and F456L positions to enhance binding affinity. Overall, this analysis pointed to a central role of F456L as a potential regulator of epistatic couplings but also suggests a potential role of structurally proximal L452 and Y453 sites supporting the improved binding interfacial interactions in the KP.3 variant. Structure-based mutational scanning of the RBD binding interfaces with different classes of RBD antibodies characterized the role of specific mutations in eliciting broad resistance to neutralization against distinct epitope classes. The results demonstrated that JN.1, KP.2 and KP.3 variants harboring the L455SS, F456L and Q493E mutations can significantly impair the neutralizing activity of RBD class-1 monoclonal antibodies, also revealing that Y453, L455 and F456 emerged as major escape hotspots for these variants. These results are consistent with the experimental data showing that antibody evasion drives the convergent evolution of L455F/S and F456L, while the epistatic interactions mediated by F456L can facilitate the favorable contribution of Q493E to restore ACE2 binding. The results support the notion that evolution of Omicron variants may favor emergence of lineages with beneficial combinations of mutations involving mediators of epistatic couplings that control balance of high ACE2 affinity and immune evasion. Our study provided support to a mechanism in which convergent Omicron mutations can promote high transmissibility and antigenicity of the virus by controlling the interplay between the binding to the host receptor and robust immune evasion profile. This may potentially be a continuous common strategy of Omicron evolution that would result in combinatorial exploration of convergent mutations to evade neutralizing antibodies and while still maintaining ACE2 binding affinity.

## Supplementary Materials

The following supporting information can be downloaded at: www.mdpi.com/xxx/s1, Figure S1: Structural alignment of the AF2-predicted RBD conformational ensemble using shallow MSA depth approach for the BA.2.86 RBD-ACE2 complex, JN.1 RBD-ACE2 complex, KP.2 RBD-ACE2 complex and KP.3 RBD-ACE2 complex. Figure S2: The distance fluctuations stability index profiles of the RBD residues obtained from MD simulations of the BA.2.86 RBD-ACE2 complex, JN.1 RBD-ACE2 complex, KP.2 RBD-ACE2 complex and KP.3 RBD-ACE2 complex. Figure S3: Structural overview of the binding interface for the S-RBD-ACE2 complexes of the Omicron XBB lineages. Figure S4: Ensemble-based mutational scanning of binding for the key RBD positions in backgrounds of the BA.2.86 variant based on MD simulations of the cryo-EM structure of the BA.2.86 RBD-ACE2 complex (pdb id 8QSQ). Figure S5: MM-GBSA binding free energy decomposition contribution of K31 and H34 ACE2 residues in BA.2.86, JN.1, BA.2.86+F456L, BA.2.86+Q493E, BA.2.86+F456L/Q493E, KP.2, BA.2.86+L455S/Q493E and KP.3 (RBD-ACE2 complexes.

## Author Contributions

Conceptualization, G.V.; Methodology, N.R., M.A., G.C., G.V.; Software, N.R., M.A., G.G., G.V.; Validation, N.R., G.V.; Formal analysis, N.R., G.V., M.A., G.G.; Investigation, N.R., G.V.; Resources, N.R., G.V., M.A, G.V.; Data curation, N.R., M.A., G.C., G.V.; Writing—original draft preparation, N.R., G.V.; Writing—review and editing, G.V.; Visualization, N.R., G.V. Supervision G.V. Project administration, G.V.; Funding acquisition, G.V. All authors have read and agreed to the published version of the manuscript.

## Funding

This research was funded by Kay Family Foundation. grant number A20-0032 and National Institutes of Health under Award 1R01AI181600-01 and Subaward 6069-SC24-11 to G.V.

## Conflicts of Interest

The authors declare no conflict of interest. The funders had no role in the design of the study; in the collection, analyses, or interpretation of data; in the writing of the manuscript; or in the decision to publish the results.

## Data Availability Statement

Data is fully contained within the article. Crystal structures were obtained and downloaded from the Protein Data Bank (http://www.rcsb.org). All simulations were performed using NAMD 2.13 package that was obtained from website https://www.ks.uiuc.edu/Development/Download/. All simulations were performed using the all-atom additive CHARMM36 protein force field that can be obtained from http://mackerell.umaryland.edu/charmm_ff.shtml. The rendering of protein structures was done with interactive visualization program UCSF ChimeraX package (https://www.rbvi.ucsf.edu/chimerax/) and Pymol (https://pymol.org/2/). All the data obtained in this work are freely available at ZENODO website link: https://zenodo.org/records/12676345.

## Acknowledgments

The authors acknowledge support from Schmid College of Science and Technology at Chapman University for providing computing resources at the Keck Center for Science and Engineering.

## Notes

### Competing Interest Statement

The authors have declared no competing interest.

