## Supplemental Figures S1-S5 for "Atomistic Prediction of Structures, Conformational Ensembles and Binding Energetics for the SARS-CoV-2 Spike JN.1, KP.2 and KP.3 Variants Using AlphaFold2 and Molecular Dynamics Simulations: Mutational Profiling and Binding Free Energy Analysis Reveal Epistatic Hotspots of the ACE2 Affinity and Immu"

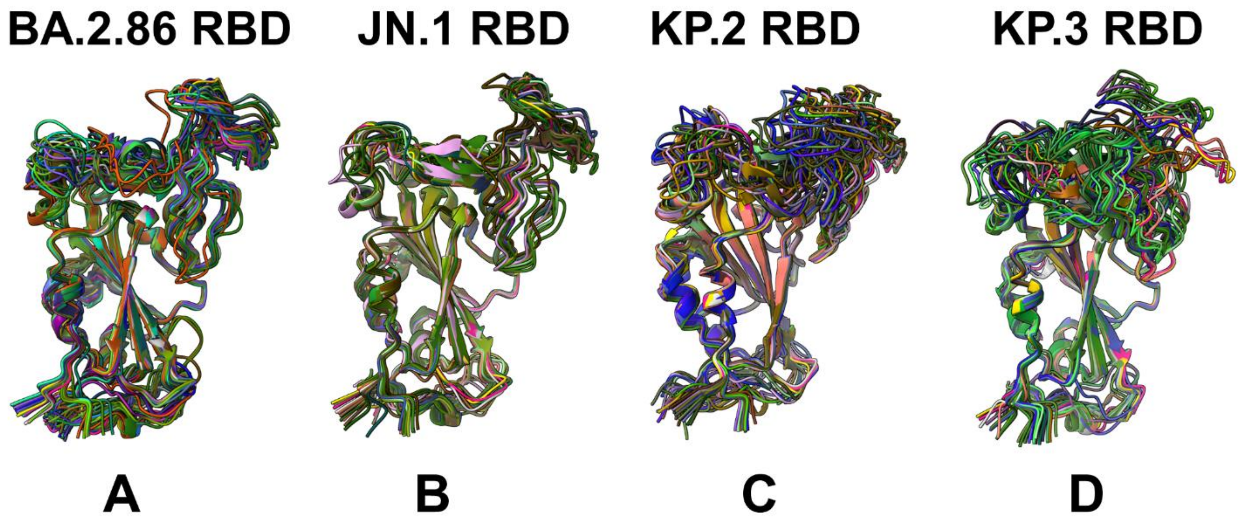


**Figure S1.** Structural alignment of the AF2-predicted RBD conformational ensemble using shallow MSA depth approach for the BA.2.86 RBD-ACE2 complex (A), JN.1 RBD-ACE2 complex (B), KP.2 RBD-ACE2 complex(C) and KP.3 RBD-ACE2 complex (D).

.

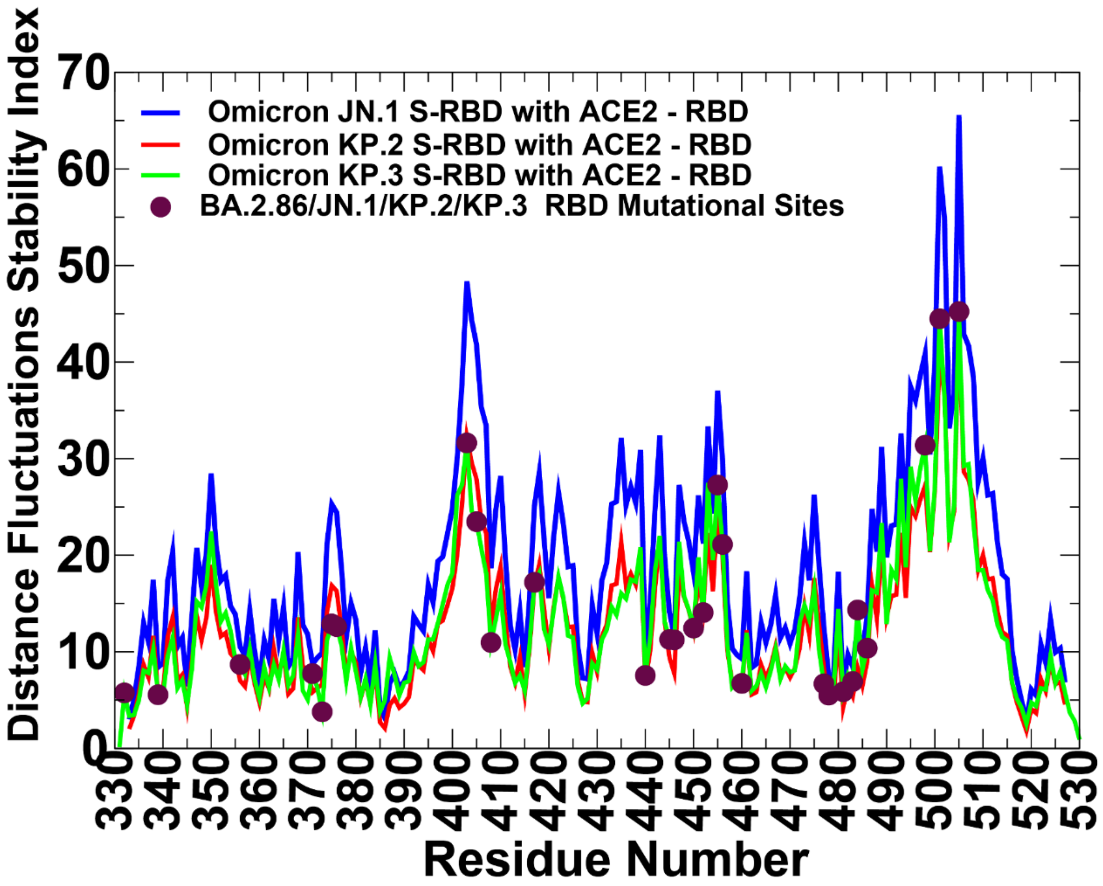


**Figure S2**. The distance fluctuations stability index profiles of the RBD residues obtained from MD simulations of the BA.2.86 RBD-ACE2 complex, JN.1 RBD-ACE2 complex, KP.2 RBD-ACE2 complex and KP.3 RBD-ACE2 complex. The profiles are shown for JN1 RBD (in blue lines) KP.2 RBD (in red lines), an KP3 RBD (in green lines) The positions of the BA.2.86/JN.1/KP.2/KP.3 mutations are highlighted in maroon colored filled circles.

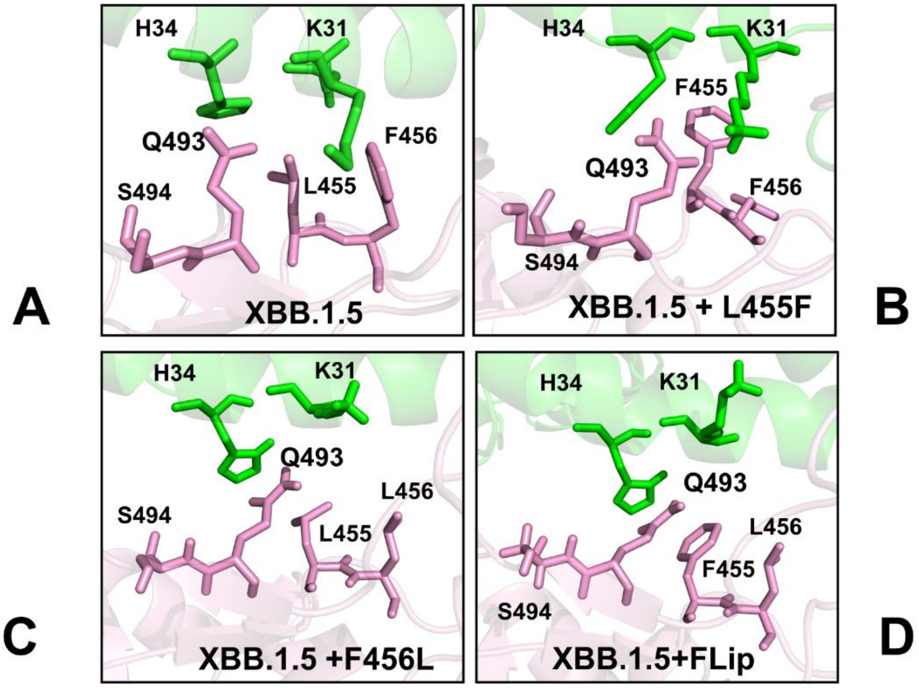


**Figure S3.** Structural overview of the binding interface for the S-RBD-ACE2 complexes of the Omicron XBB lineages. (A) A closeup of the binding interface residues Q493, L455 and F456 in the cryo-EM structure of the XBB.1.5 RBD-ACE2 complex, pdb id 8WRL. The RBD residues are in cyan sticks, the ACE2 residues H3 and K31 are in green sticks. (B) A closeup of the binding interface residues Q493, L455F and F456 in the AF2-predicted best model of the XBB.1.5+L455F complex with ACE2. (C) A closeup of the binding interface residues Q493, L455 and F456L in the cryo-EM structure of the XBB.1.5+F456L complex with ACE2, pdb id 8WTD (C) A closeup of the binding interface residues Q493, L455F and F456L in the cryo-EM structure of the XBB.1.5+L455F/F456L FLip complex with ACE2, pdb id 8WRH. The binding interface residues are shown in pink sticks for RBD sites and green sticks for ACE2 sites. The RBD interface residues Q493, L455F and F456L undergo rearrangements in the XBB.1.5+FLip complex.

c

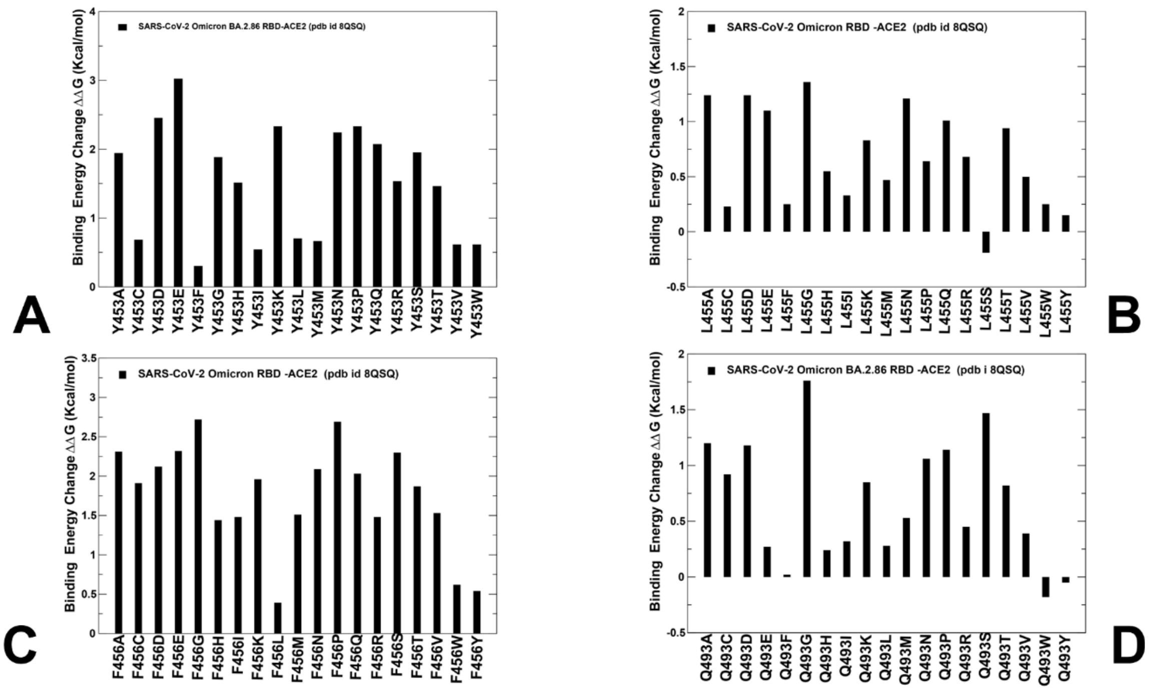
A
SARS-CoV-2 Omicron BA.2.86 RBD-ACE2 (pdb id 8QSQ)
Binding Energy Change ΔΔG (Kcal/mol)
Y453A Y453C Y453D Y453E Y453F Y453G Y453H Y453I Y453K Y453L Y453M Y453N Y453P Y453Q Y453R Y453S Y453T Y453V Y453W
B
SARS-CoV-2 Omicron RBD -ACE2 (pdb id 8QSQ)
Binding Energy Change ΔΔG (Kcal/mol)
L455A L455C L455D L455E L455F L455G L455H L455I L455K L455M L455N L455P L455Q L455R L455S L455T L455V L455W L455Y
C
SARS-CoV-2 Omicron RBD -ACE2 (pdb id 8QSQ)
Binding Energy Change ΔΔG (Kcal/mol)
F456A F456C F456D F456E F456G F456H F456I F456K F456L F456M F456N F456P F456Q F456R F456S F456T F456V F456W F456Y
D
SARS-CoV-2 Omicron BA.2.86 RBD -ACE2 (pdb i 8QSQ)
Binding Energy Change ΔΔG (Kcal/mol)
Q493A Q493C Q493D Q493E Q493F Q493G Q493H Q493I Q493K Q493L Q493M Q493N Q493P Q493R Q493S Q493T Q493V Q493W Q493Y

**Figure S4.** Ensemble-based mutational scanning of binding for the key RBD positions in backgrounds of the BA.2.86 variant based on MD simulations of the cryo-EM structure of the BA.2.86 RBD-ACE2 complex (pdb id 8QSQ). The profiles of computed binding free energy changes ΔΔG upon 19 single substitutions for Y453 residue in BA.2.86 (A), L455 in BA.2.86 (B), F456 in BA.2.86 (C), and Q493 in BA.2.86 (D. The binding free energy changes are computed using conformational ensembles obtained from MD simulations. The binding free energy changes are shown in black-colored filled bars. The positive binding free energy values ΔΔG correspond to destabilizing changes and negative binding free energy changes are associated with stabilizing changes.

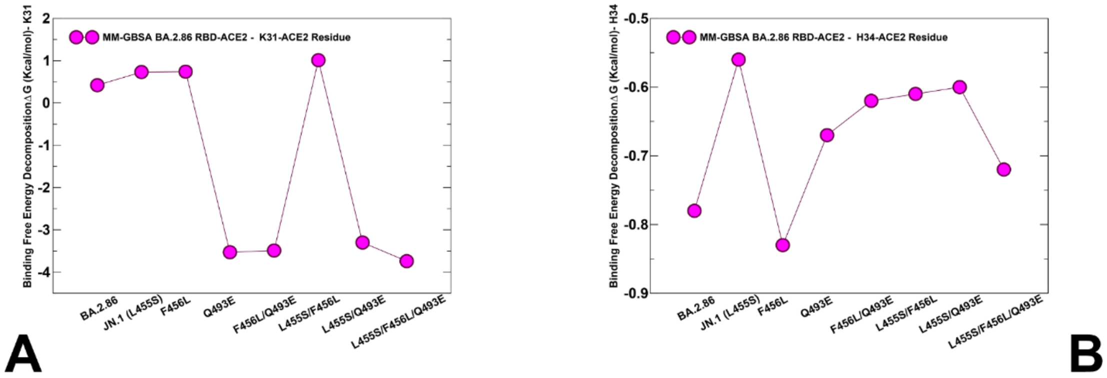


**Figure S5.** MM-GBSA binding free energy decomposition contribution of K31 and H34 ACE2 residues in BA.2.86, JN.1, BA.2.86+F456L, BA.2.86+Q493E, BA.2.86+F456L/Q493E, KP.2, BA.2.86+L455S/Q493E and KP.3 (RBD-ACE2 complexes. (A) The MM-GBSA decomposition contribution of the binding energies for K31-ACE2 residue. (B) The MM-GBSA decomposition contribution of the binding energies for H34-ACE2 residue.
